# Clarithromycin Exerts an Antibiofilm Effect against *Salmonella typhimurium* rdar Biofilm Formation, and Transforms the Physiology towards an Apparent Oxygen-depleted Energy and Carbon Metabolism

**DOI:** 10.1101/2020.05.02.068536

**Authors:** Munira Zafar, Humera Jahan, Sulman Shafeeq, Manfred Nimtz, Lothar Jänsch, Ute Römling, M. Iqbal Choudhary

## Abstract

Upon biofilm formation, production of extracellular matrix components and alteration in physiology and metabolism allows bacteria to build up multicellular communities which can facilitate nutrient acquisition during unfavorable conditions and provide protection towards various forms of environmental stresses to individual cells. Thus, bacterial cells become tolerant against antimicrobials and the immune system within biofilms. In the current study, we evaluated the antibiofilm activity of the macrolides clarithromycin and azithromycin. Clarithromycin showed antibiofilm activity against rdar (red, dry and rough) biofilm formation of the gastrointestinal pathogen *Salmonella typhimurium* ATCC14028 Nal^r^ at 1.56 μM subinhibitory concentration in standing culture and dissolved cell aggregates at 15 μM in a microaerophilic environment suggesting that the oxygen level affects the activity of the drug. Treatment with clarithromycin significantly decreased transcription and production of the rdar biofilm activator CsgD, with biofilm genes such as *csgB* and *adrA* to be consistently downregulated. While *fliA* and other flagellar regulon genes were upregulated, apparent motility was downregulated. RNA sequencing showed a holistic cell response upon clarithromycin exposure, whereby not only genes involved in the biofilm-related regulatory pathways, but also genes that likely contribute to intrinsic antimicrobial resistance, and the heat shock stress response were differentially regulated. Most significantly, clarithromycin exposure shifts the cells towards an apparent oxygen- and energy-depleted status, whereby the metabolism that channels into oxidative phosphorylation is downregulated, and energy gain by degradation of propane 1,2-diol, ethanolamine and L-arginine catabolism is upregulated. This initial analysis will allow the subsequent identification of novel intrinsic antimicrobial resistance determinants.

## INTRODUCTION

Elevated tolerance against antibiotics is mainly promoted by biofilm formation of the organism (1,2). Biofilms are multicellular communities of microorganisms embedded into the self-produced extracellular matrix. Biofilm formation occurs on abiotic as well as biotic surfaces, but the multicellular behavior is also displayed as cell aggregation (free-floating in liquid, so-called flocks) and pellicle formation (3,4). Among the various functions of bacterial biofilms, provision of nutrients and to defend their replicating entities during unfavorable environmental conditions are among major characteristics. As biofilms are consequently tolerant against disinfectants, antimicrobials and the immune system of the host, according to the NIH (National Institute of Health), up to 80% of human bacterial infections are associated with biofilm formation (5,6). Alongside microbial cells, up to 90% of the biofilm mass can be comprised of the extracellular matrix, which can consist of self-produced exopolysaccharide, proteins, lipids and extracellular DNA (eDNA), but also environmental or host components. The extracellular matrix can be extensively hydrated, differ in solubility, and its component cross-link thus providing the physical- and chemical shield towards antimicrobials and other stress conditions (7).

The Gram-negative food borne pathogen *Salmonella typhimurium*, characteristically forms biofilms which contributes to the perseverance of *Salmonella* in the food industry (8). Foodborne diseases arise due to the contamination of food-products such as poultry, but also vegetables and fruits by *Salmonella*. According to the United States Centers for Disease Control and Prevention (CDC), almost 48 million cases of foodborne diseases occur annually and sequentially leading to 128,000 hospitalizations and 3000 deaths (9).

*S. typhimurium* displays a specific colony morphology, which is characterized by red, dry and rough appearance on agar plates containing Congo red. This biofilm, termed as rdar morphotype, expresses the exopolysacharide cellulose and proteinaceous curli as biofilm matrix components (10). Biofilm development is controlled by the key transcriptional regulator CsgD (curli subunit gene D), a LuxR family protein which activates transcription of the *csgBAC* operon encoding curli fiber components, and indirectly governs the activity of the cellulose synthase by promoting transcription of the diguanylate cyclase AdrA (11,12,2). CsgD plays a pivotal role in switching between biofilm, motility and virulence state(s) of bacteria, regulated by various, defined and yet undefined, stimuli, which alter the production of the second messenger cyclic di-GMP (13,14,15) Antibiotics have a short, not more than 85 years history to be used systematically in the medical setups to treat and eradicate bacterial infections, yet the emergence of drug resistant and multidrug resistant organisms has been frequently detected early on. As biofilm formation leads to chronic infections refractory to be treated by antimicrobial therapy, identifying a role of antibiotics in modulating biofilm formation can lead to a rational approach in the development of successful anti-biofilm therapies. Evaluating FDA approved drugs for their antibiofilm activity can reduce development time and cost up to 40% (16).

Macrolides, a class of broad-spectrum antibiotics, were the third class of antibiotics to be discovered used since 1952 in clinical practice. Erythromycin, the first discovered natural macrolide, has thereby been replaced by the more effective second generation semi-synthetic macrolides clindamycin and azithromycin (17). Macrolides selectively inhibit protein biosynthesis by binding to the 23S RNA of the 50 S subunit of the ribosome, halting translation in bacteria mainly by blocking the entrance tunnel for the nascent peptide chain (18,19,20,21). An antibiofilm role of macrolides has been observed in various species of bacteria, but the underlying mechanisms have not been resolved (22). Apart from their role as antibiotics, macrolides possess immunomodulatory and anti-inflammatory properties (23).

The aim of this study was to identify the antibiofilm potential of different established antibiotics, and to unravel the underlying regulation. We found that the macrolide antibiotic clarithromycin possesses antibiofilm activity against rdar biofilm forming *S. typhimurium* UMR1. RNA sequencing showed that clarithromycin at subinhibitory concentrations, besides differential expression of biofilm-related genes such as the central activator *csgD*, causes a distinct multiplex physiological response, which will lead to the characterization of antibiofim and antimicrobial targets and the design of novel combinatorial antimicrobial strategies. Such does upregulation of genes and proteins related to ribosome functionality and the heat shock response indicate compensatory innate resistance mechanisms against the protein synthesis inhibiting 23S RNA targeting drug clarithromycin. Furthermore, the bacterial cell experiences apparent oxygen and carbon depletion upon clarithromycin treatment as indicated by the upregulation of genes encoding the trimethylamine-N-oxide reductase and involved in anaerobic propane-1,2-diol and L-arginine degradation, and downregulation of genes involved in pathways channeling into oxidative phosphorylation such as the Krebs, glyoxylate and 2-methylcitrate cycle.

## MATERIALS AND METHODS

### 96-well Antibiofilm Assay

The bacterial strains used in the study were wild type *S. typhimurium* UMR1 (ATCC14028 Nal^r^; positive for rdar biofilm formation at 28 °C, and *S. typhimurium* MAE50 (UMR1 Δ*csgD*) as rdar biofilm negative control. Strains were grown on Luria broth (LB) without salt agar plates containing Congo red (40μg/mL) and Coomassie brilliant blue (20 μg/mL). Single colonies of bacteria were inoculated in 5 mL LB and grown overnight at 37 °C with shaking. The cell density was adjusted to 0.1 OD_600_ and resuspended in LB without salt broth, suspended into a 96-well plate with 2-fold serial dilutions of the drug starting from 100 μM and incubated for 48 h at 28 °C to allow the formation of biofilm. Biofilm was washed, and adherent cells were stained with 0.2% crystal violet solution. After dissolution of the dye, OD_600_ was measured to assess biofilm formation, and % biofilm inhibition was calculated using the following formula to evaluate anti-biofilm activity:

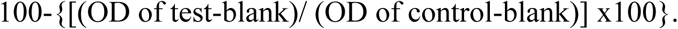

Growth inhibition was assessed by transferring 100 μL of the cell suspension from the 96-well plate into a fresh plate and the OD_600_ of the cell suspension was measured. Growth inhibition was calculated with the same formula as mentioned above.

### Biofilm formation in liquid culture

Cell aggregation was observed upon *S. typhimurium* growth under microaerophilic condition in liquid culture. Overnight cultures of *S. typhimurium* were inoculated to OD_600_ 0.02 with 60 % of the volume of the flask filled with LB without salt medium and incubated at 28 °C with 150 rpm shaking for 16 h. After 16 h of incubation, cell aggregation was visually analysed and documented, and upon resuspension the OD_600_ of the cell suspension was measured.

### Assessment of cell aggregation by light microscopy

In order to observe the effect of the drug on cell aggregation and morphology, light microscopy was performed. 20 μL of cell suspension from each flask was spread onto a glass slide, and cell aggregates were observed under the light microscope.

### Detection of Protein Expression by Western Blotting

To monitor the expression level of the major biofilm regulator CsgD Western blot analysis was carried out. *S. typhimurium* UMR1 cells with different concentrations of clarithromycin (3.25, 7.5, 15, 30 and 60 μM), cells without treatment as positive control, and *S. typhimurium* MAE50 as negative control were grown under microaerophillic conditions with shaking at 150 rpm at 28 °C. After 16 h, the cell suspension was centrifuged and 2.5 mg of pelleted cells dissolved in 100 μl sample buffer. Commassie brilliant blue staining of sodium dodecyl sulfate-polyacrylamide gel electrophoresis (4% stacking and 12% resolving gel) was carried out to analyse the protein content. Equal amounts of protein were separated, and transferred to a polyvinylidene difluoride membrane (Immobilon P; Millipore) using the semidry Western blotter Mini-Sub Cell GT from Bio-Rad. Detection of CsgD was carried out using a polyclonal anti-CsgD peptide antibody (1:2000) and horseradish peroxidase-conjugated goat anti-rabbit IgG (1:3000). CsgD was detected by LUMI-Light kit from Roche.

### RNA Isolation

The RNA from biofilm-forming bacteria was isolated as previously described (11). *S. typhimurium* UMR1 untreated and treated with 15 μM clarithromycin was grown under microaerophilic conditions with shaking at 150 rpm at 28 °C for 16 h. Cells were harvested by centrifugation and disrupted by lysis buffer (50 mM Tris HCL pH 8, 8% sucrose, 0.5% Triton, 20mM EDTA, 4mg/ mL), 200 μL acidic phenol was added, and samples were then heated at 65 °C for 15 min, while shaking. The sample was cooled on ice for 5 min and centrifuged at 13000 rpm. The aqueous phase was collected and extracted once with an acid phenol:chloroform mix, and twice with chloroform. Finally, RNA was precipitated with 3 volumes of 100% ethanol and 0.3 M sodium acetate overnight at -80 °C. Pellets were washed with 1 mL 70% ethanol, and air-dried after the supernatant was removed. Pellet was dissolved in 100 μL DEPC-treated distilled water. After DNAse (Ambion RiboPure-Yeast DNase) treatment, the RNA samples were analysed by denaturating gel electrophoresis to assess DNA contamination and integrity of the isolated RNA. If DNA contamination was still present, the RNA was repeatedly incubated with DNAse. The RNA concentration was measured by Nano drop and the quality of RNA preparation assessed by Agilent Bioanalyzer 2100. The DNA-free RNA was stored at -80 °C until use.

### cDNA synthesis and quantitative RT PCR

cDNA was prepared from 1 μg of isolated DNA free RNA using the high-capacity cDNA Reverse Transcription Kit (Applied Biosystems) and the reaction was run in thermal cycler according to the protocol. For quantitative RT-PCR, reaction volume was 25 μl, 3 μM forward and reverse primer was prepared, 5 μl cDNA (1:20 diluted), 10 μl SYBR green (iTaq Universal SYBR Green Supermix, Bio-Rad) filled up with 3 μl DEPC-treated water was added in a 96-well plate and sealed with adhesive film. Before running qPCR, the plate was spun down for 3 min at 3000 rpm at room temperature. Quantitative real-time PCR was performed using the 7500 Real-Time PCR system (Applied Biosystems). Relative transcript abundance was determined by the 2ΔΔCT method using the 7500 SDS Software v1.3.1 (Applied Biosystems) with the *rpsV* gene coding for the 30 S ribosomal subunit protein S22 and the *recA* gene used for internal normalization. Primers for quantitative real-time RT-PCR are listed in Supplementary Table-1.

### RNA Sequencing and Data processing

RNA samples were depleted from the ribosomal RNAs and subsequently sequenced on a Nextseq 550 with medium output flowcell 150 cycles (2×75 PE). The resulting bcl files were converted and demultiplexed to fastq using the bcl2fastq program. STAR was used to index the *S. typhimurium* strain 14028S genome (NCBI Reference Sequence: NC_016856.1), and to subsequently align the fastq files. Mapped reads were then counted in annotated genes using feature Counts. The annotations were also obtained from NCBI in gff3 format. The count table from feature Counts was imported into R/Bioconductor, and differential gene expression was performed using the EdgeR package and its general linear models pipeline. For the gene expression analysis, genes that had 1 count per million in 3 or more samples were used. Dispersion was tagwise calculated and TMM was used to normalize the samples. Genes were termed significant if they had an FDR adjusted *p* value under 0.05. A Volcano and Scatter plot was drawn with Exel indicating all differentially expressed genes with a *p*-value <0.01.

### Mass spectrometry for Protein identification

Differential protein bands were cut out from the SDS-PAGE gel, subjected to in-gel trypsin digestion and extracted and purified by standard procedure (extraction by H_2_O, trifluoroacetic acid and acetonitrile and analyzed by UltiMate 3000 RSLC nano LC system connected to an Orbitrap Velos mass spectrometer (Thermo Scientific). MS survey scans were alternated with MS/MS scans of daughter ion spectra obtained from peptide components by collision induced dissociation (CID). All survey spectra were recorded using the orbitrap detector with a resolution of 60 k and the daughter ion spectra of peptide components were recorded using the ion trap. The experimental setup was as follows: the peptide mixture was loaded onto a 75 μm × 2 cm precolumn (Acclaim PepMap 100, C18, 3 μm, Thermo Scientific) and eluted over 60 min using a 75 μm × 25 cm analytical column (Acclaim PepMap RSLC, C18, 2 μm, Thermo Scientific) with a gradient buffer 0–95% acetonitrile in 0.1% formic acid. The data were acquired using the Xcalibur 2.3 software (Thermo Scientific) and processed with the Peaks 6 software (Bioinformatics Solution). MS/MS data were searched versus the 208140 entry *Salmonella enterica* database extracted from the NCBI non-redundant database. Search parameters were fixed modification carbamidomethylation (57.02) and methionine oxidation (15.99) as variable modification; cleavage by trypsin; parent mass error tolerance ±10.0 ppm and fragment mass error tolerance 0.3 Da. Maximal missed cleavage of 1 and significance threshold p < 0.05. Selected identified proteins from individual bands are reported.

## RESULTS

### Clarithromycin, but not azithromycin shows predominantly an antibiofilm effect

*S. typhimurium* UMR1 cells were inoculated at an OD_600_ 0.1 and incubated with serially 2-fold diluted drug from 0.02 μM to 100 μM. In order to assess the growth inhibition, the OD_600_ of the cell suspension was estimated. To analyze the antibiofilm potential of clarithromycin, we assessed biofilm formation by staining the bacterial cells adherent to the wall of well with crystal violet. An antibiofilm effect was judged to be present when the percentage of biofilm inhibition was consistently and significantly exceeding the percentage of growth inhibition at a distinct concentration of clarithromycin. Among the two screened macrolides, azithromycin and clarithromycin, only clarithromycin showed an antibiofilm activity (Figure-1, -2).

**Figure-1:**
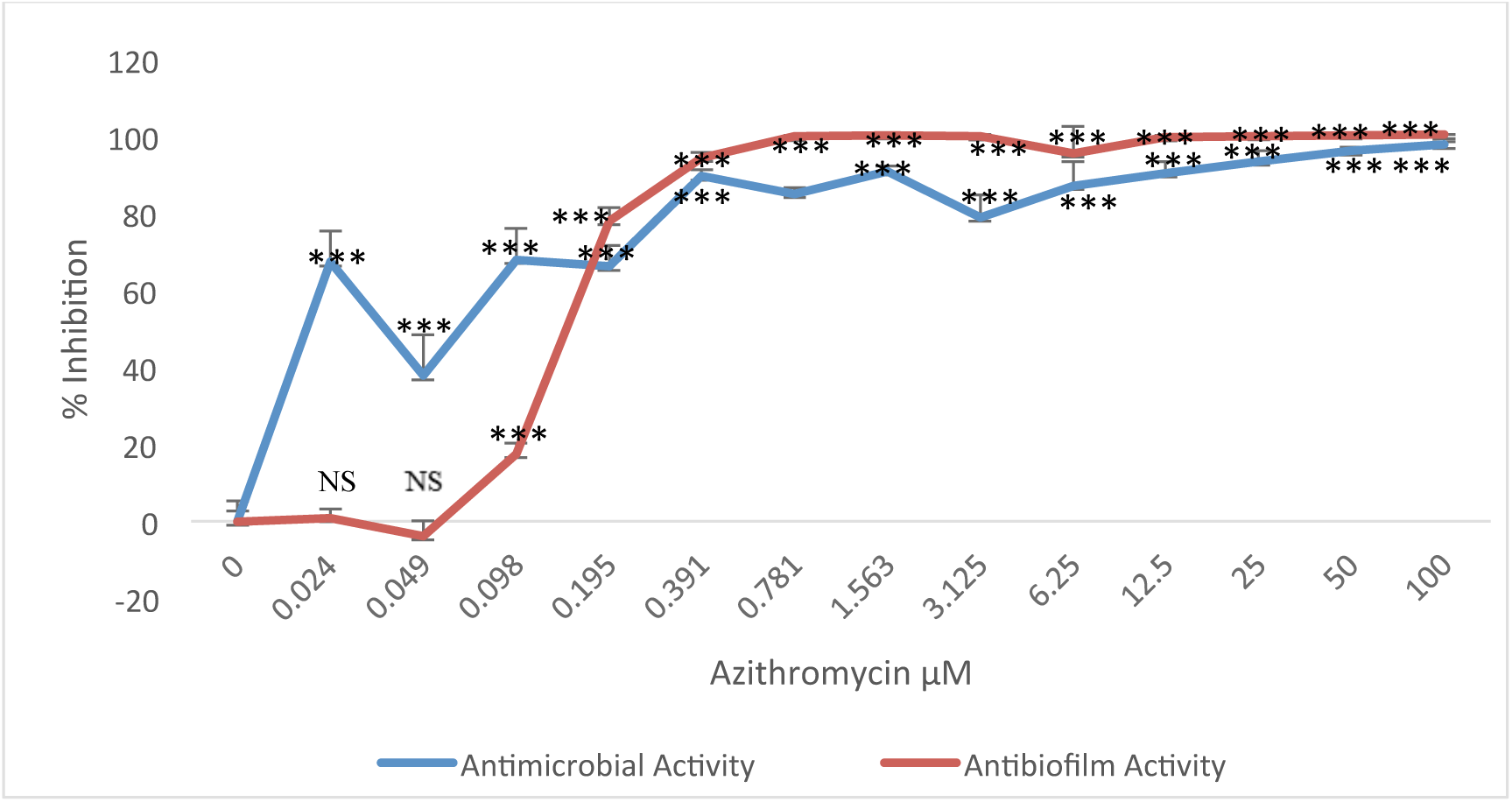
Antibiofilm activity and growth inhibition of azithromycin against *S. typhimurium* UMR1 as assessed by staining the biofilm after 96-well plate incubation with crystal violet and measurement of the OD_600_ of the cell suspension, respectively.. Azithromycin pre-dominantly showed an antimicrobial effect. The data shown are mean ± SEM of 6 technical replicates.

**Figure-2:**
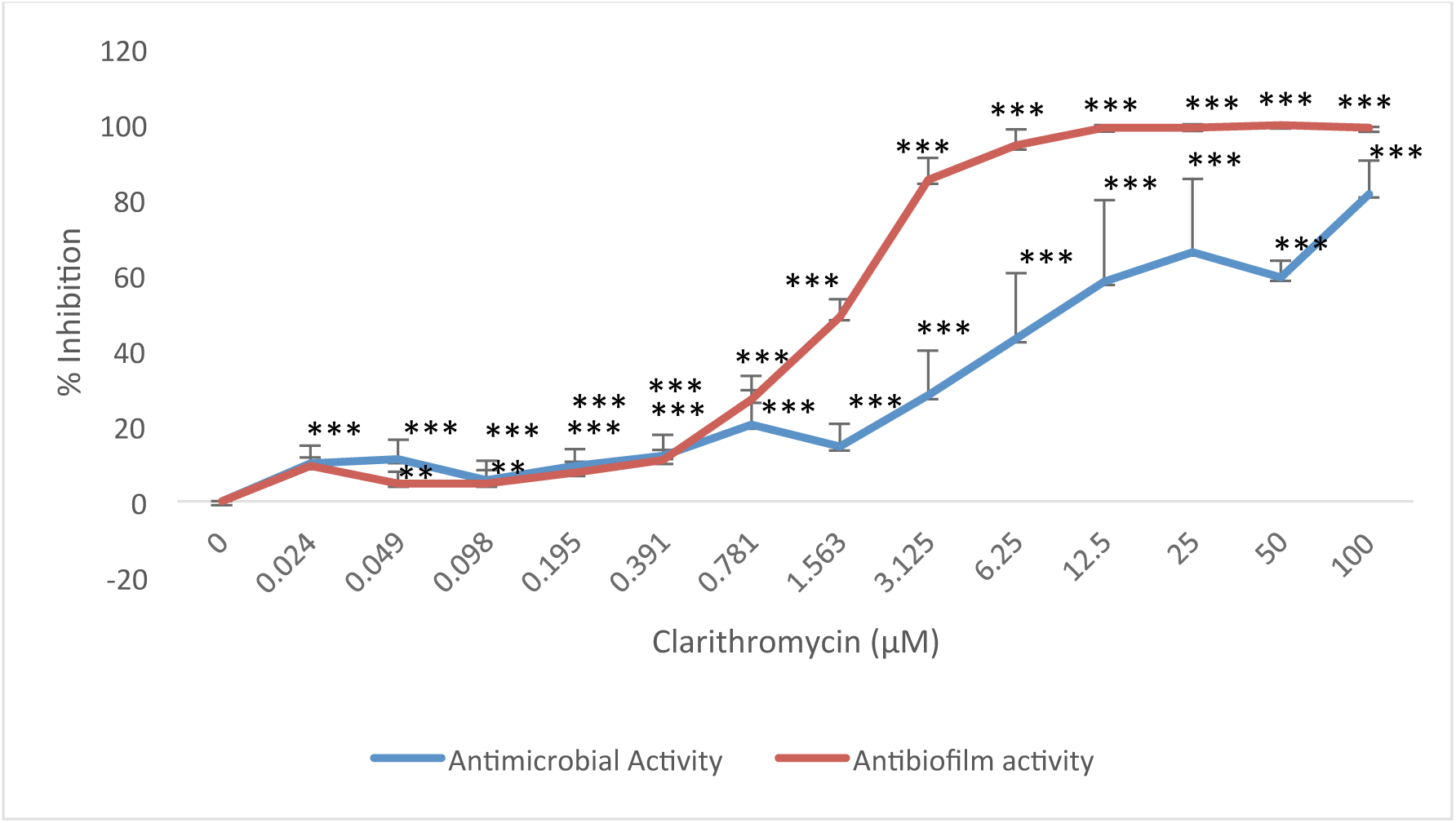
Antibiofilm activity and growth inhibition of clarithromycin against *S. typhimurium* UMR1 as assessed by staining the biofilm after 96-well plate incubation with crystal violet and measurement of the OD_600_ of the cell suspension, respectively. At 1.56 μM, clarithromycin predominantly showed antibiofilm activity and conseqeuntly a less pronounced growth inhibition. Untreated *S. typhimurium* UMR1 was the positive control and *S. typhimurium* MAE50 was the negative control. Data shown are the mean ± SEM of 2 biological with 6 technical replicates each.

### Antibiofilm effect of Clarithromycin

We observed that at 1.56 μM clarithromycin, the biofilm was inhibited by 49 %, whereas the antimicrobial activity was 14 %, showing that clarithromycin has an antibiofilm activity at this particular concentration as seen in Figure-2.

### Clarithromycin prevents Cell Aggregation in Liquid Culture

To investigate the antibiofilm effect of clarithromycin in another biofilm model, the cells were grown under microaerophilic conditions in liquid culture. Under these experimental conditions, biofilm formation of *S. typhimurium* is also expressed as macroscopically visible cell aggregates (Figure-3).

**Figure-3:**
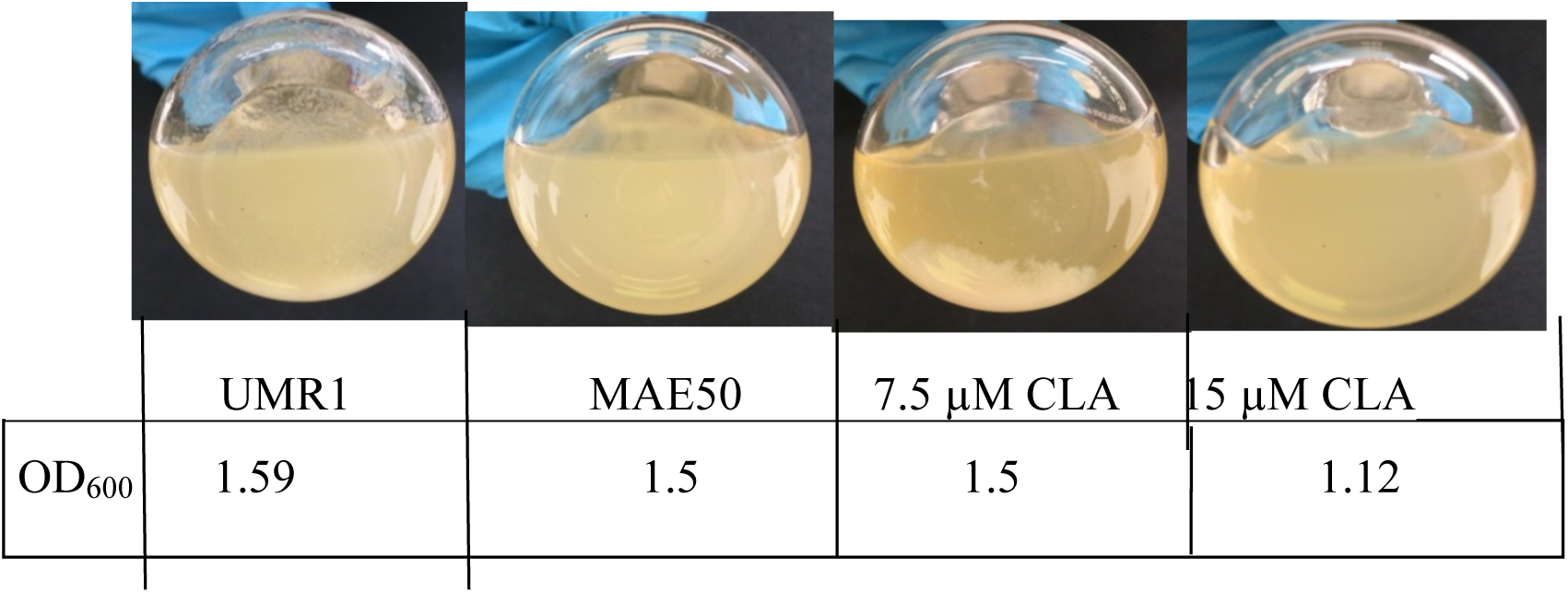
Effect of clarithromycin (CLA) on cell aggregation of *S. typhimurium* UMR1 in liquid culture. *S. typhimurium*. UMR1 was grown under microaerophillic conditions in liquid culture at 28° for 16 hours and 150 rpm shaking, where biofilm formation expressed in the form of cell aggregates. Cell aggregation is visible in wild type *S. typhimurium* UMR1 untreated and up to 7.5 μM. Upon treatment with 15 μM clarithromycin, no cell aggregation was observed. To evaluate the growth inhibitory effect of clarithromycin, OD_600_ was estimated. Minor growth inhibition compared to the untreated *S. typhimurium* UMR1 was observed upon treatment with 15 μM clarithromycin.

After inoculation of *S. typhimurium* with different concentrations (3.25, 7.5, 15, and 30 μM) of clarithromycin, cells aggregates were clearly visible in considerable amounts in *S. typhimurium* UMR1, as described previously (23). In contrast, at 7.5 μM clarithromycin, still substantial clumps were visible, whereas at 15 μM clarithromycin, cell aggregation was not observed and only planktonic cells were found to be present (Figure-3).

### Microscopic analysis

The amount of cell aggregation and aggregate morphology was assessed by bright field microscopy at the different concentrations (3.25, 7.5, 15, and 30 μM) of clarithromycin. Large cell aggregates with a tight consistency were observed for *S. typhimurium* UMR1 (Figure-4). Upon treatment with clarithromycin, cell aggregation was found to be concentration dependent decreased. Already at 3.25 μM, a pronounced effect of clarithromycin could be noticed as distorted cell aggregates with less defined edges and loosely attached clumps were present. At μM clarithromycin, cell aggregates significantly decreased, and consisted of smaller and more loosely connected clumps. At 15 μM clarithromycin, only planktonic cells were visible demonstrating the gradual inhibition of biofilm formation of *S. typhimurium* UMR1 upon clarithromycin treatment.

**Figure-4:**
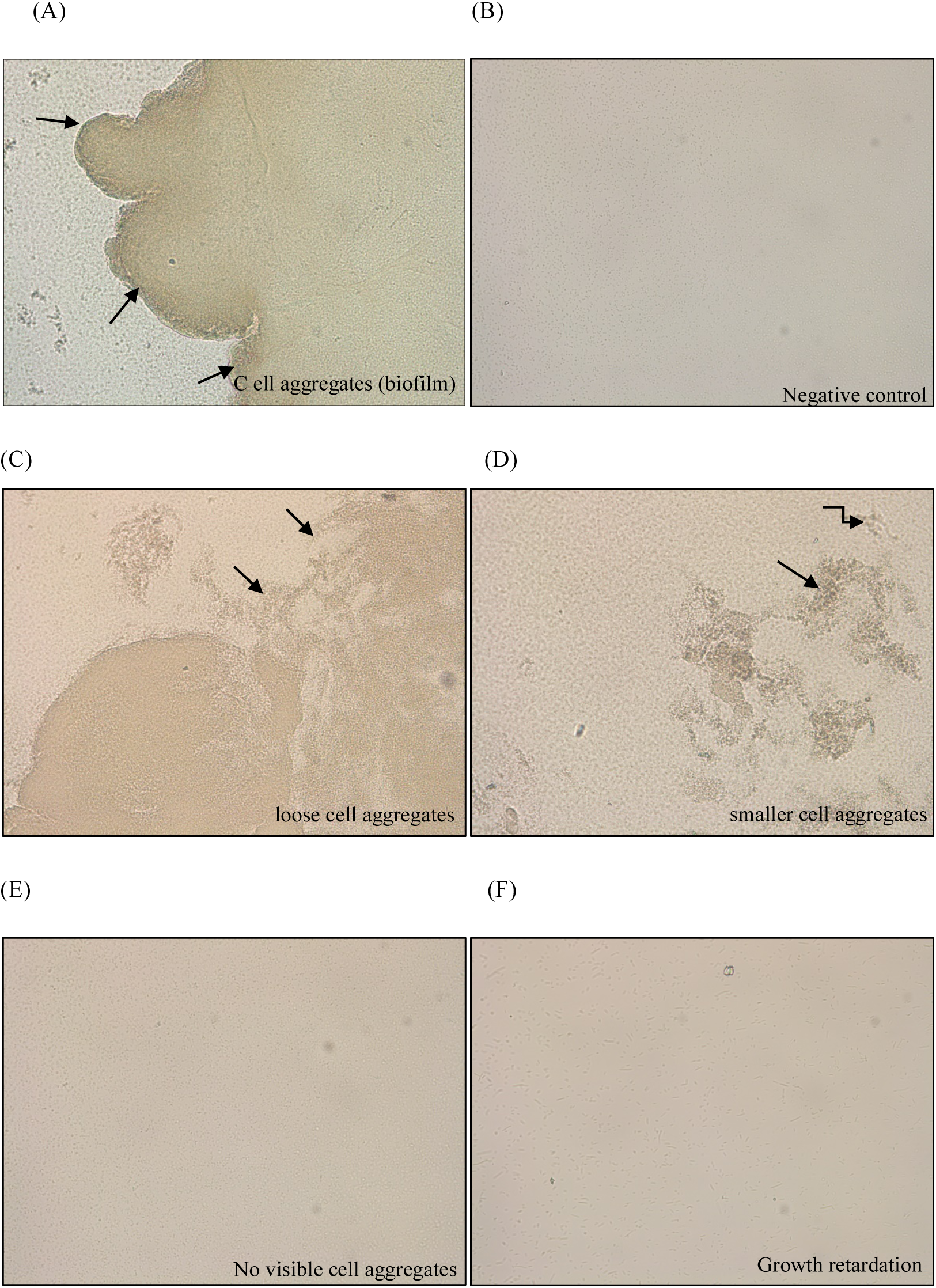
Light microscopy images of cell aggregates of (A) *S. typhimurium* UMR1; (B) *S. typhimurium* MAE50 (negative control) and *S. typhimurium* UMR1 treated with 3.25 μM (C), 7.5 μM (D), 15 μM (E) 30 μM (F) clarithromycin. A representative part of the cell suspension was spread on a glass slide and cell aggregation was documented. Arrows point to cell aggregates. All images were taken at 20X magnification.

### Expression of the Biofilm Regulator CsgD is decreased upon Treatment with Clarithromycin

CsgD is the central transcriptional activator required for rdar biofilm formation (3,111,6). Consequently, we monitored the expression of CsgD upon exposure to clarithromycin (Figure 5). Upon treatment of *S. typhimurium* UMR1 under microaerophilic conditions in liquid culture with 3.25, 7.5, 15, and 30 μM of clarithromycin, a gradual decrease in the expression levels of CsgD was detected by Western blot analysis. Consistent with the cell aggregation phenotype, at 15 μM clarithromycin, almost no CsgD protein production was detected (Figure-5). Therefore, the underlying mechanism of clarithromycin to inhibit rdar biofilm formation is consequently to target the regulatory pathway(s) leading to *csgD* expression.

**Figure-5:**
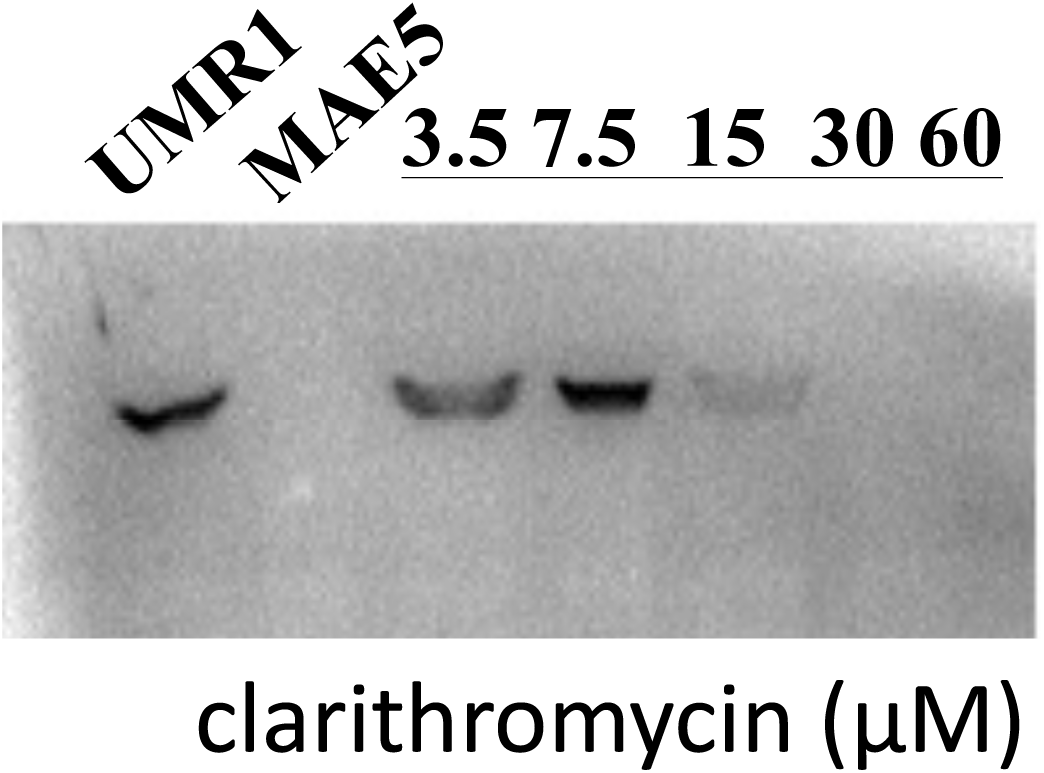
CsgD expression in liquid culture under microaerophilic growth. Cells were harvested from the *S. typhimurium* UMR1 culture without, and with different concentrations (3.25, 7.5, 15, 30, 60 μM) of clarithromycin after 16 hours of incubation at 28°C and 150 rpm shaking after. The commassie stained gel is provided in the supplementary figure-1. The CsgD expression level was significantly diminished upon treatment with 15 μM clarithromycin.

### Differential Expression of Biofilm and Motility Genes upon Treatment with Clarithromycin as assessed by qRT-PCR

As CsgD protein expression was downregulated upon treatment with clarithromycin, we estimated biofilm gene expression by quantitative real time RT-PCR in order assess on which level clarithromycin interferes with biofilm formation (Figure-6). The expression level of *csgD*, and consequently the expression level of *csgB*, coding for the minor subunit of curli fimbriae, and *adrA*, encoding the diguanylate cyclase required for activation of the extracellular matrix component cellulose, was found to be decreased by 50%, as compared to the untreated control. In addition, expression of *bcsA*, coding for the cellulose synthase, was decreased, although *bcsA* expression is not regulated by *csgD* (25). On the other hand, expression of *fliA* encoding the sigma factor 28 of the flagella regulon and *fliB* coding for flagellin N-methylase, was increased. This differential expression pattern holds, although the *rpsV* and *recA* reference genes were both subsequently found to be downregulated by RNA sequencing.

**Figure-6:**
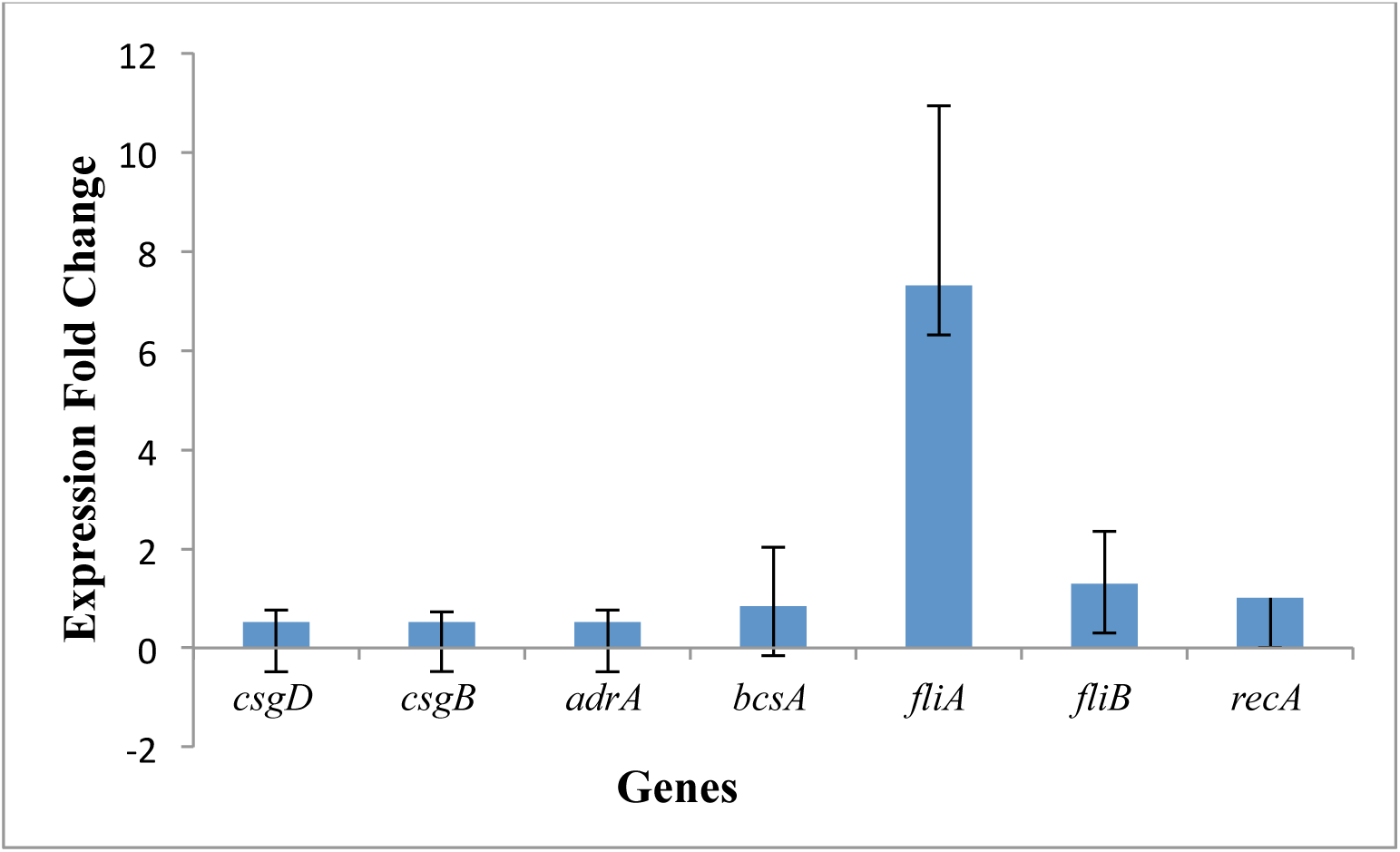
Expression of target genes *(csgD, csgB, adrA, bcsA, fliA, fliB*) at 15 μM clarithromycin. While, *csgD, csgB, adrA* and *bcsA* were found down regulated after treatment with clarithromycin,*fliA* and *fliB* were upregulated. The fold-expression upon treatment versus untreated control are indicated. For RNA isolation, cells were harvested after 16 hours of incubation under microaerophilic condition at 28°C and 150 rpm shaking. The data were expressed as mean ± SEM of two experiments.

### Clarithromycin treatment significantly alters the transcriptome of rdar biofilm forming *S. typhimurium* UMR1

Treatment with clarithromycin down regulated the expression of the major biofilm regulator *csgD*. In order to assess differential expression of the regulatory pathway(s) leading to the inhibition of rdar biofilm formation upon treatment with clarithromycin, as well as to unravel additional alterations of the cellular transcriptome upon clarithromycin treatment, we performed RNA sequencing (Figure-7). *S. typhimurium* UMR1 grown under microaerophilic conditions, 16 h at 28°C and 150 rpm shaking, was treated with 15 μM clarithromycin. Of the 4586 annotated genes of *S. typhimurium* ATCC14028S, 353 genes were >4-fold, while 107 genes were >8-fold differentially regulated. Supplementary tables 2 and 3 display the 30 most up- and down-regulated genes.

**Figure-7:**
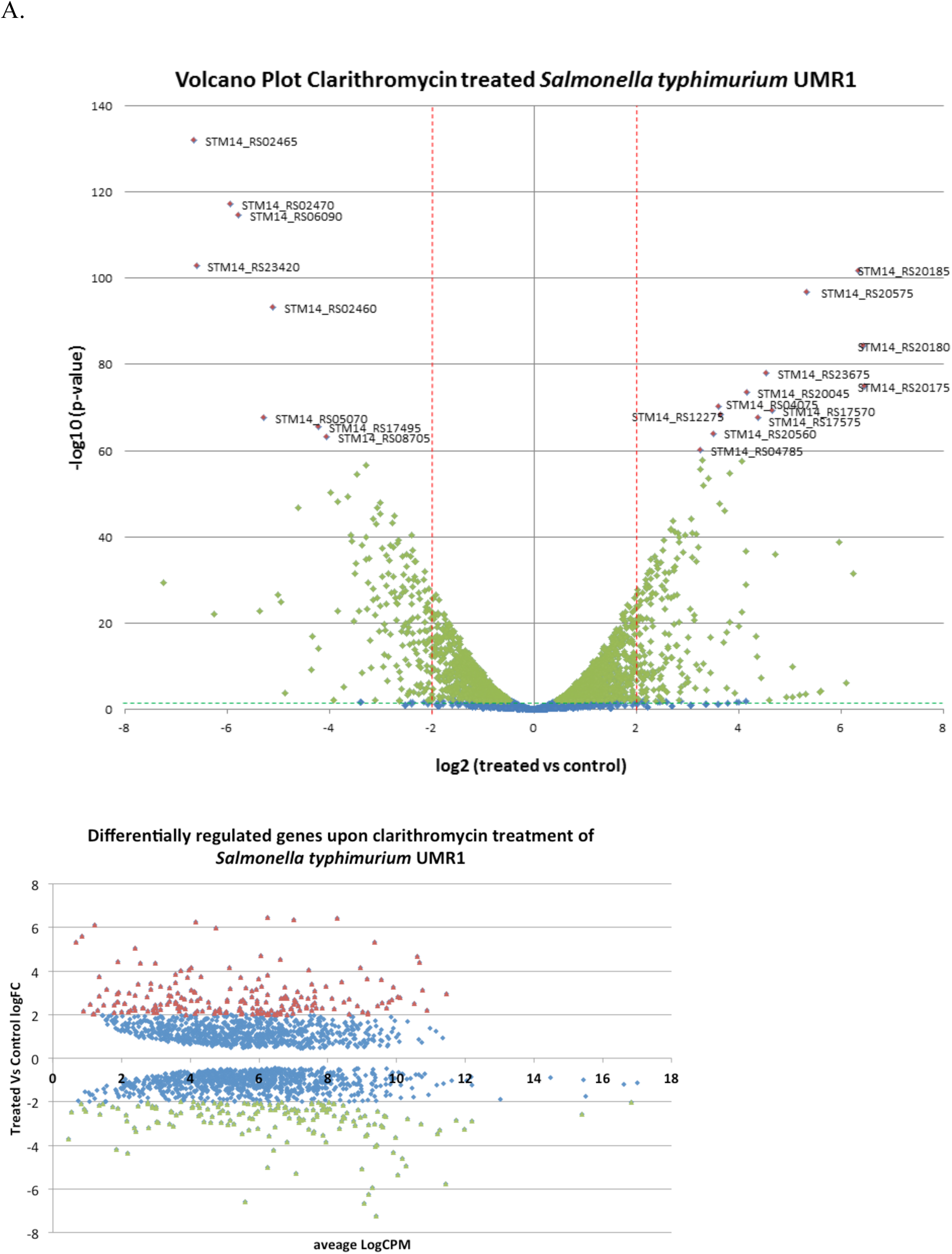
Genes differentially regulated upon treatment of *S. typhimurium* UMR1 with 15 μM clarithromycin grown under microaerophilic conditions. (A) Volcano plot indicating the significantly differentially regulated genes. Dotted red line indicates genes differentially regulated more than 4-fold. Dotted green line indicates p>0.01 threshold of statistical significance. Red symbol, 20 genes most significantly differentially regulated more than 4-fold. Green symbol, significantly differentially regulated genes with p<0.01 threshold. Blue symbol, genes differentially regulated and/or not significantly regulated (threshold p>0.01). (B) Scatter blot indicating the expression level of significantly differentially regulated genes.

Consistent with the observed rdar biofilm phenotype and the previously performed qRT-PCR experiments, the transcriptional data showed that genes of the divergently transcribed *csgDEFG* and *csgBAC* operons were consistently downregulated (Supplementary Table-4), although those genes were not most downregulated. In addition, other genes involved in biofilm formation are downregulated, such as the gene coding for the cyclic di-GMP synthase AdrA that is required for cellulose expression. Upregulation of the gene for the truncated EAL protein STM0551, a post-transcriptional repressor, indicates downregulation of type 1 fimbriae (26) Motility and biofilm formation are two fundamentally opposite life styles of bacteria (27). In contrast to biofilm genes, most flagellar regulon genes were upregulated including *flhD* and *flhC* encoding the class 1 central flagella regulator FlhD_4_C_2_ and the class 2 gene *fliA* encoding the flagellar sigma 28 factor, although a few class 2 and class 3 genes including *fliC* and the phosphodiesterase *yhjH* were downregulated (Supplementary Table-5). The dramatic downregulation of three paralogous *rpoS* regulated genes coding for <60 amino acid KGG rich intrinsically disordered proteins (YmdF, YciG and STM14 RS08420) required for swimming and swarming motility (28), however, might indicate their decisiveness for the observed lack in swimming motility (data not shown).

Among the most downregulated genes (Supplementary Table-3) were two ferritin-like genes involved in the storage of iron in non-toxic form equally as the ferritin-like protein Dps (29,30). Clarithromycin-exposed cells might be depleted of the osmoprotectant trehalose, as genes for biosynthesis and break-down enzymes are downregulated (24,31). Furthermore, consistently downregulated are genes that encode Krebs-cycle enzymes, including associated pathways such as the 2-methylcitrate cycle, the GABA shunt, and the glyoxylate cyclase. Indeed, downregulation of the genes coding for propionate-CoA ligase and formate C-acetyl transferase indicates that also the production of substrates channeling into these pathways is blocked.

The most upregulated genes upon treatment with clarithromycin were the three genes of the *torCAD* operon encoding a cytochrome C-type protein, the trimethylamine N-oxide (TMAO) reductase, and its chaperone (Supplementary Table-2) (32). Trimethylamine N-oxide serves as an alternative electron acceptor in anaerobic respiration. Furthermore, genes coding for the propane-1,2-diol microcompartment and, partially, ethanolamine utilization pathway were both upregulated, although their parallel upregulation has been reported to be mutually exclusive (33,34). Propane-1,2-diol and ethanolamine are usually utilized as carbon and energy source under anaerobic conditions supported by the alternative electron acceptor tetrathionate (34). Another pathway highly transcriptionally upregulated are genes involved in the degradation of L-arginine under anaerobiosis (35). In addition, the genes of the pathway(s) leading to biosynthesis of branched-chain amino acids, leucine, isoleucine, and valine are consistently upregulated (36).

The macrolide clarithromycin targets the ribosome by binding to the 23S RNA within the 50S ribosomal subunit to selectively inhibit protein synthesis, to prevent assembly of ribosomes, and to trigger tRNA dissociation. We found distinct genes involved in protein synthesis and homeostasis to be upregulated. For example, genes coding for ribosomal proteins such as for the 50S ribosomal proteins L21 and L27 and rRNA and ribosomal protein modifiying genes such as the genes for 23S rRNA (guanine(745)-N(1))-methyltransferase and the 30S ribosomal protein S5 alanine-N-acetyltransferase, respectively, were more than 16-fold upregulated (Supplementary Table-6). Furthermore, distinct t-RNAs and t-RNA modification genes were also upregulated. In particular, four t-RNAs for Arginine are upregulated >4-fold (with one Arginine t-RNA downregulated). Supplementary Table-6 shows the heatmap for differentially regulated ribosome associated genes. As also the genes encoding the trigger factor and the heat shock response, including genes encoding the heat shock sigma factor RpoH and holding chaperons IbpA and IbpB were upregulated, we conclude that the clarithromycin exposed cell is under severe proteotoxic stress. A highly specific response is therefore set up to overcome the effect of clarithromycin on prevention of protein synthesis and peptidyl t-RNA dissociation in order to ensure proper protein translation, folding and to prevent protein aggregation.

The upregulation of the gene for the multidrug efflux MFS transporter MdtM indicates that the cell actively pumps out the macrolide through MdtM. The downregulation of the open reading frame of the outer membrane porin OmpC might indicate a first line innate immune response to protect against the macrolide.

Furthermore, we preliminary investigated how the alterations on the transcriptional level translate into changes on the protein level upon treatment with clarithromycin (Figure-8). One-dimensional protein gels showed a substantial change in the protein expression profile, but no significant protein degradation up to 60 μM clarithromycin (Figure-8). In congruence with the transcriptome data, potentially upregulated proteins upon exposure to clarithromycin included ribosomal proteins, while potentially downregulated proteins included the ferritin-like metalbinding protein YciE, and proteins that belong to major metabolic pathways such as the Krebs cycle. Two-dimensional gel elecrophoresis or gel-free approaches such as SILAC (stable isotope labelling by amino acid in cell culture) is required to verify the identity of upregulated proteins. Figure-9 summarizes the overall physiological consequences of the differentially regulated genes on treatment of *S. typhimurium* rdar biofilm (37) with clarithromycin.

**Figure 8:**
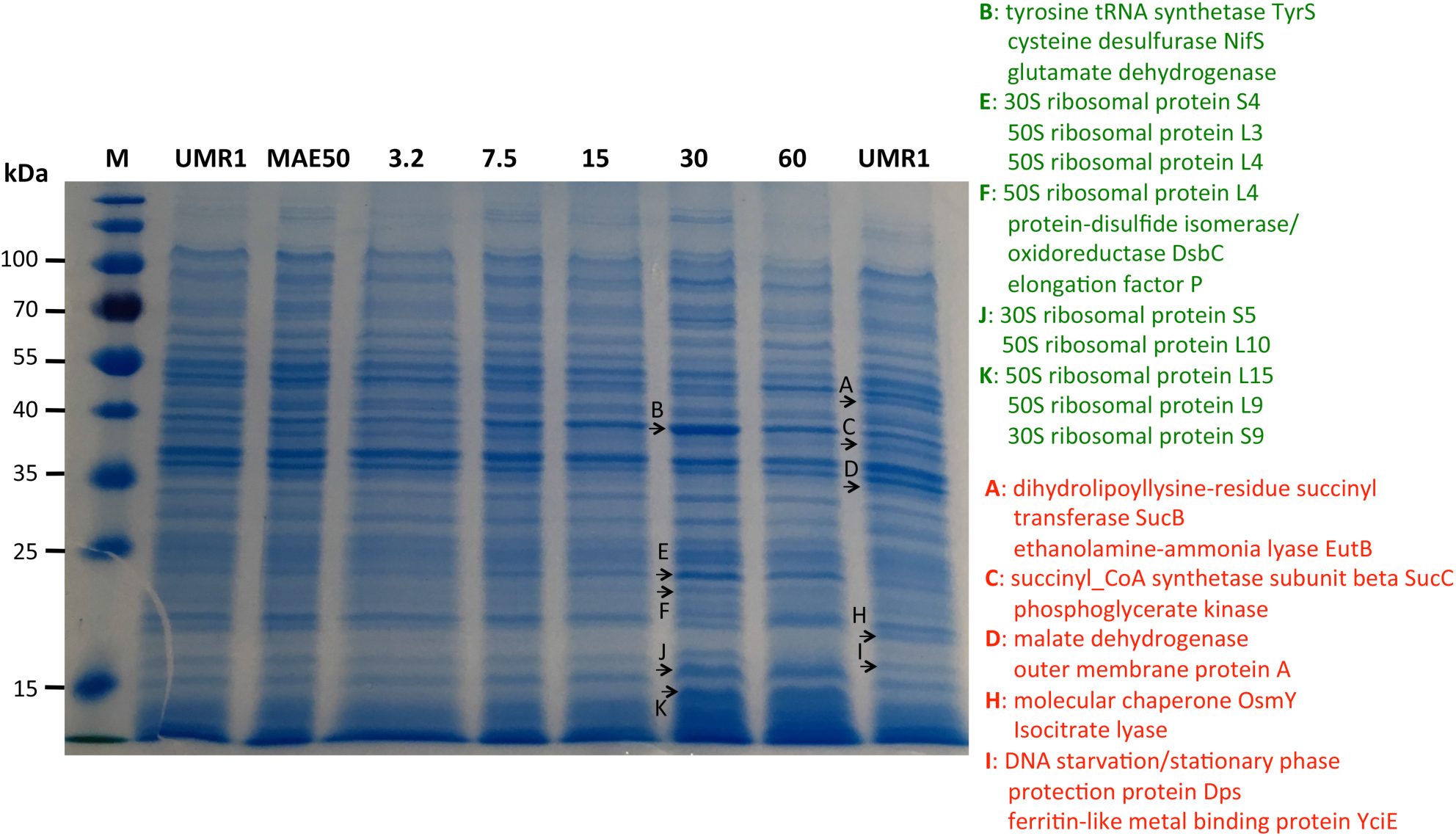
Changes in protein expression upon clarithromycin treatment.

**Figure 9:**
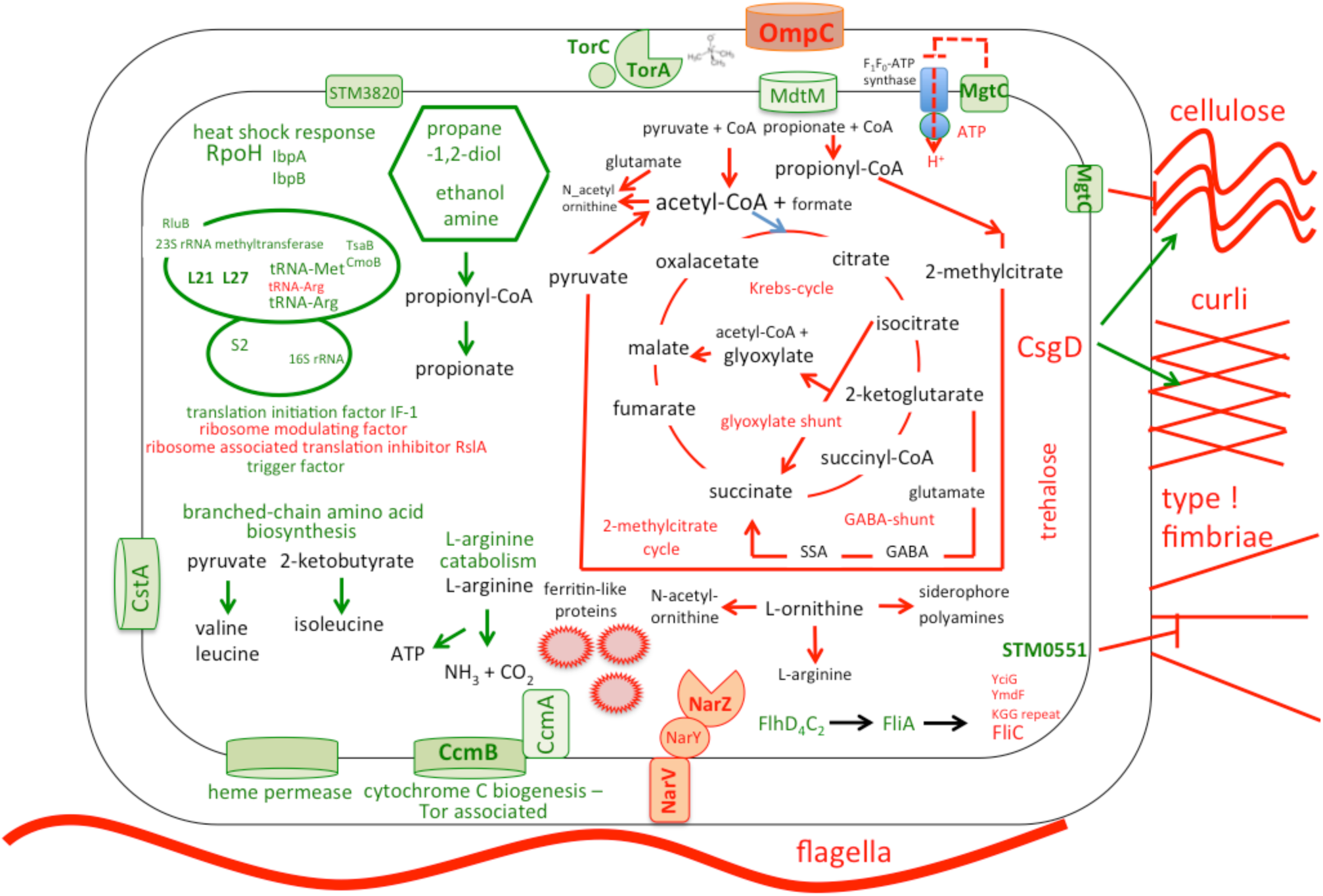
Summarizing figure of major physiological and metabolic changes in rdar biofilm forming *S. typhimurium* upon exposure to 15 μM clarithromycin. Further explanation, see text.

## DISCUSSION

Biofilm formation of microorganisms impairs antimicrobial treatment strategies of chronic infections because of the tolerance of biofilms against antibiotics as well as the immune system. Biofilm formation is particularly pronounced on implanted medical devices due to the lack of effective antimicrobial defense mechanisms (38). One strategy to tackle biofilm infections is interference with Quorum sensing, as this external microbial communication mechanism can promote biofilm formation (39). Antibiotics have traditionally been used to eradicate bacterial infections in the last 85 years. Using the existing drugs to target biofilms can be time and cost effective, avoid safety tests with the side effects known (40).

Clarithromycin can target various pathogens and infections (41,42,43). Moreover a biofilm inhibitory effect for this drug (alone or in combination) against *Pseudomonas aeruginosa, Helicobacter pylori* and *Staphylococcus aureus* has been reported (44,45). We have evaluated the role of this macrolide antibiotic primarily as an antibiofilm agent against *S. typhimurium*. RNA-sequencing unraveled not only the molecular mechanisms and various pathways involved in the biofilm inhibiting effect of clarithromycin, but also demonstrated major metabolic alterations that might provide the basis to rationally design combinatorial treatment strategies.

After screening a library of well-investigated antibiotics, we observed that clarithromycin among other macrolides possessed antibiofilm properties. Surprisingly, we observed a growth condition dependent effect of clarithromycin, as 1.56 μM clarithromycin was sufficient in the 96-well standing culture biofilm assay to prevent biofilm formation, while >15 μM clarithromycin was required to prevent cell aggregation and growth under microaerophilic conditions in shaking culture. Based on our RNA-sequencing data, we conclude that clarithromycin sets the cell into an apparent state of oxygen depletion and strongly interferes with pathways channeling into the oxidative phosphorylation, which might explain the oxygen-dependent effect (Figure-9; see below).

The expression of the *csgD* biofilm activator, which is indicative for rdar biofilm formation can dramatically alter in response to environmental conditions and temperature (24). Clarithromycin treatment targets expression of *csgD* and biofilm-associated genes, which could be used in controlling infections related to *Salmonella* (46). A study in the *Escherichia coli* outbreak strain O104:H4 shows biofilm formation and virulence genes to be coexpressed suggesting an antibiofilm strategy can be an effective antivirulence strategy (47). Moreover, *csgD* and the extracellular matrix components of *Salmonella*, curli and cellulose, have been shown to play a role in persistence (48). The genes determinative for downregulation of *csgD* expression are not entirely obvious. Among the known genes positively regulating *csgD* expression, genes encoding the sigma factor RpoS and the transcriptional regulator MlrA are moderately downregulated. However, the differential regulation of small RNAs, several of which affect *csgD* expression, still needs to be analyzed in detail.

Several other effects of clarithromycin might contribute to downregulation of biofilm formation. We observed high upregulation of genes involved in the electron transport chain and anaerobic respiration. Alteration in enzymes involved in electron transport chain interferes with biofilm formation in Gram-positive and Gram-negative bacteria (49). Furthermore, there exists a correlation between multidrug efflux pumps and biofilm in *Salmonella*, as biofilm formation is compromised upon deletion or inhibition of multidrug efflux pumps (44). Our data suggests that the gene for the multidrug efflux transporter MdtM is upregulated, which can be an innate resistance mechanism as upregulation of efflux pumps confers multiple drug resistance in bacteria (50). It has been previously shown that a *csgD* deletion mutant accumulates tricarboxylic (Krebs) cycle intermediates in order to block gluconeogenesis. Furthermore, cells in biofilms have a higher ATP requirement (51). We have observed the downregulation of the Krebs cycle and related pathways upon clarithromycin treatment induced *csgD* downregulation which prevents efficient energy gain through oxidative phosphorylation, a mechanism which might contribute to inhibition of biofilm formation. In the same line, the gene for the virulence factor MgtC is upregulated. MgtC downregulates cellulose biosynthesis and inhibits the ATP synthase (52). An efficient inhibition of the ATP synthase might stimulate the cell to seek alternative sources to produce the energy equivalent ATP.

Moreover, genes related to iron metabolism were highly downregulated upon clarithromycin treatment (Supplementary Table-3). *E. coli* downregulates the iron related gene expression during optimal growth conditions to prevent stress response mediated by high levels of iron (53), although in *S. typhimurium* during multicellular swarming motility, significantly upregulates the genes for iron metabolism (54).

Surprisingly, most upregulated were genes involved in the use of alternative carbon and energy sources under anaerobic conditions. These were genes involved in anaerobic propane-1,2-diol, ethanolamine and L-arginine degradation. The high upregulation of a gene encoding the carbon starvation transporter CstA might facilitate the acquisition of carbon sources. Furthermore, genes coding for the trimethylamine N-oxide reductase were most dramatically upregulated. Cumulatively, also taking into consideration the downregulation of Krebs cycle genes and genes of associated pathways, these findings indicate that clarithromycin sets the cells under apparent oxygen and energy depletion. Of note, bactericidal, but not bacteriostatic drugs involve the Krebs cycle and the production of reactive oxygen species (55). Supporting the oxygen-depletion hypothesis, the TMAO reductase is required for survival of cells experiencing sudden oxygen depletion (56). We predict that the deletion of those highly upregulated genes will increase the sensitivity for clarithromycin, and can lead to the detection of novel components contributing to the innate resistance against clarithromycin. Indeed, the cellular response as experienced by a subinhibitory concentration of antibiotics and analyzed by RNA-sequencing has successfully identified intrinsic antimicrobial resistance components (57,58). Besides the identification of potentially novel intrinsic antimicrobial resistance factors associated with bacterial energy gain, more obvious components contributing to intrinsic antimicrobial resistance are associated with the ribosome, protein biosynthesis and quality control, direct targets and consequences of clarithromycin exposure. However, only distinct genes are substantially upregulated suggesting a highly specific response. Among those genes is the *rrmA* gene coding for the 23S_rRNA_(guanine(745)-N(1))-methyltransferase that has been shown to aid elongation of the polypeptide chain (59), several 50 and 30S ribosomal proteins and ribosome modulating enzymes (60,61,62) in addition to tRNAs and tRNA modifying enzymes (63). Of note, although ribosome-associated, none of these factors has previously been identified to contribute to clarithromycin resistance, which includes identified resistance mechanisms such as methylation of the 23S rRNA at adenosine2058 and ribosomal protein L4 and L22 overexpression (64). Furthermore, genes involved in initiation of translation, protein folding and quality control are upregulated. In summary, RNA sequencing revealed a highly specific response against clarithromycin which extended beyond the inhibition of rdar biofilm formation and upregulation of intrinsic antimicrobial resistance components.

## ACKNOWLEDGEMENT

We thank the Bioinformatics and Expression Analysis core facility at the Karolinska Institutet, Sweden for performing RNA-sequencing and data processing. This research was funded by Swedish Research Links. We also acknowledge the financial support of the Pakistan Academy of Sciences (Grant No. PAS/1439).

## Supplemental data

**Supplemental-Table 1.**
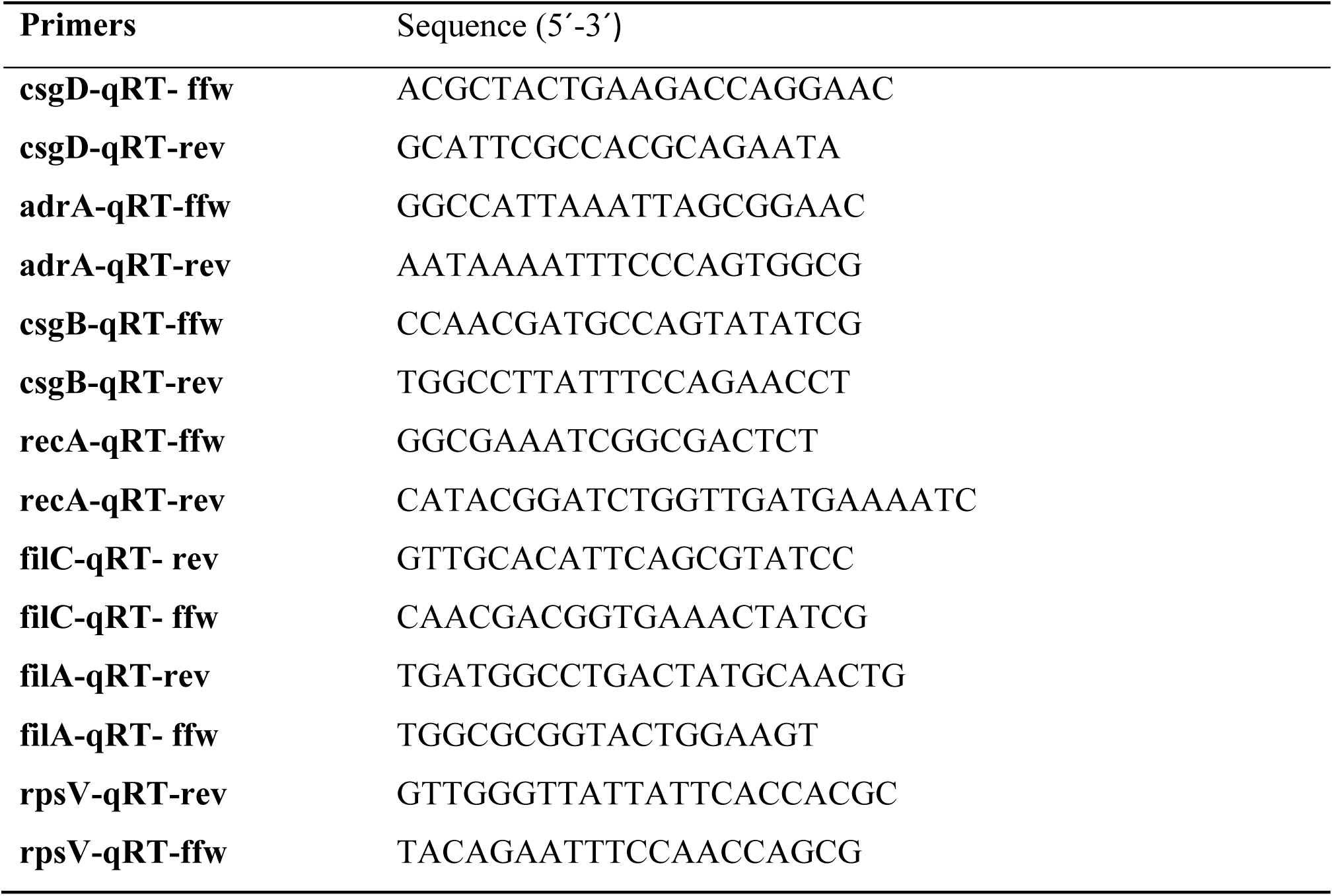
Primers used for the real time PCR.

**Supplemental-table 2.** 30 most downregulated genes upon treatment with 15 μM clarithromycin.

| S. No | Gene ID | Product | Log 2<br>(Treated<br>Vs<br>Control) | P Value |
| --- | --- | --- | --- | --- |
| 1. | STM14 RS09550 | Ferritin like domain containing protein YciE | -7.23448 | 2.39E-29 |
| 2. | STM14 RS02465 | Bifunctional 2-methylcitrate<br>dehydratase/aconitate hydratase | -6.64236 | 3.1E-132 |
| 3. | STM14 RS23420 | Ornithine carbamoyl transferase | -6.59155 | 8.97E-104 |
| 4. | STM14 RS09545 | Ferritin like domain containing protein YciF | -6.25568 | 3.019E-22 |
| 5. | STM14 RS02470 | Propionate CoA ligase | -5.92532 | 7.26E-117 |
| 6. | STM14 RS06090 | Small intrinsically disordered KGG protein<br>YmdF | -5.76585 | 9.03E-115 |
| 7. | STM14 RS08420 | Small intrinsically disordered KGG protein<br>GsiB | -5.34933 | 5.706E-23 |
| 8. | STM14 RS05070 | Arginine ABC transporter substrate-binding<br>protein | -5.27504 | 6.906E-68 |
| 9. | STM14 RS02460 | 2-methylcitrate synthase | -5.09532 | 4.47E-93 |
| 10. | STM14 RS07160 | Four-helix bundle copper binding protein | -5.00141 | 6.81E-27 |
| 11. | STM14 RS09540 | Small intrinsically disordered KGG protein<br>YciG | -4.93784 | 4.59E-25 |
| 12. | STM14 RS09940 | Hypothetical small cytoplasmic protein MisL | -4.60275 | 8.58E-47 |
| 13. | STM14 RS07165 | tRNA-Arg | -4.35863 | 7.22E-10 |
| 14. | STM14 RS09555 | Manganese catalase family protein CotJC | -4.32319 | 1.98E-17 |
| 15. | STM14 RS17495 | Argininosuccinate synthase | -4.21714 | 3.47E-66 |
| 16. | STM14 RS06855 | Alpha/beta hydrolase | -4.2027 | 3.40E-15 |
| 17. | STM14 RS08705 | Alcohol dehydrogenase AdhP | -4.04233 | 2.77E-63 |
| 18. | STM14 RS21980 | Isocitrate lyase | -3.97071 | 4.35E-51 |
| 19. | STM14 RS03680 | Glutathione-dependent formaldehyde dehydrogenase | -3.8326 | 3.32E-23 |
| 20. | STM14 RS21985 | Bifunctional isocitrate dehydrogenase kinase/phosphatase | -3.83036 | 4.43E-49 |
| 21. | STM14 RS11145 | Hypothetical phage-related protein | -3.71789 | 4.78E-06 |
| 22. | STM14 RS05315 | Formate C-acetyl transferase | -3.63444 | 7.08E-50 |
| 23. | STM14 RS21670 | N-acetyl-gamma glutamyl phosphate reductase | -3.56888 | 1.24E-40 |
| 24. | STM14 RS23685 | NAD-dependent succinate semialdehyde dehydrogenase | -3.5645 | 1.47E-39 |
| 25. | STM14 RS08665 | Malto oligosyltrehalose synthase | -3.52295 | 5.17E-21 |
| 26. | STM14 RS21975 | Malate synthase A | -3.48992 | 8.877E-32 |
| 27. | STM14 RS12550 | Porin OmpC | -3.48192 | 1.27E-34 |
| 28. | STM14 RS21675 | Acetyl glutamate kinase | -3.46071 | 6.31E-56 |
| 29. | STM14 RS01125 | Glucose uptake inhibitor SgrT | -3.42052 | 2.03E-23 |
| 30. | STM14 RS08760 | Nitrate reductase subunit alpha | -3.35489 | 6.39E-39 |

**Supplemental-table 3.** 30 most upregulated genes upon treatment with 15 μM clarithromycin.

| S. No | Gene ID | Product | Log 2<br>(Treated<br>Vs<br>Control) | P- Value |
| --- | --- | --- | --- | --- |
| 1. | STM14 RS20175 | Molecular chaperone Tor D | 6.47168 | 2.96E-75 |
| 2. | STM14 RS20180 | Trimethylamine N-oxide reductase Tor A | 6.44235 | 3.55E-84 |
| 3. | STM14 RS20185 | Pentaheme c-type cytochrome Tor C | 6.34787 | 2.86E-103 |
| 4. | STM14 RS11400 | Propanediol utilization microcompartment protein PduB | 6.24118 | 2.78E-32 |
| 5. | STM14 RS25015 | *Salmonella* restricted hypothetical protein | 6.12656 | 9.92E-07 |
| 6. | STM14 RS23410 | Arginine deiminase | 5.97575 | 1.62E-39 |
| 7. | STM14 RS12475 | Heme ABC transporter permease | 5.61585 | 4.84E-05 |
| 8. | STM14 RS20575 | Acetolactate synthase 2 catalytic subunit | 5.33777 | 2.77E-96 |
| 9. | STM14 RS03365 | truncated EAL protein STM0551 | 5.33358 | 2.28E-04 |
| 10. | STM14 RS11415 | Propanediol dehydratase small subunit | 5.05466 | 1.43E-10 |
| 11. | STM14 RS23750 | Carbon starvation protein | 4.72518 | 9.57E-37 |
| 12. | STM14 RS17570 | 50S ribosomal protein L27 | 4.65689 | 1.05E-68 |
| 13. | STM14 RS23675 | Multidrug efflux MFS transporter MdtM | 4.55136 | 3.23E-79 |
| 14. | STM14_RS11435 | Propanediol utilization microcompartment protein PduK | 4.44997 | 4.19E-08 |
| 15. | STM14_RS17575 | 50S ribosomal protein L21 | 4.39303 | 2.01E-67 |
| 16. | STM14_RS11395 | Propanediol utilization microcompartment protein PduA | 4.37134 | 6.23E-13 |
| 17. | STM14_RS11410 | Propanediol dehydratase medium subunit PduD | 4.35164 | 1.14E-17 |
| 18. | STM14_RS20045 | Acetolactate synthase large subunit | 4.15821 | 1.42E-73 |
| 19. | STM14_RS19885 | MgtC family protein | 4.15602 | 2.00E-29 |
| 20. | STM14_RS20580 | Acetolactate synthase 2 small subunit | 4.15189 | 8.41E-37 |
| 21. | STM14_RS20615 | Keto acid_reductoisomerase | 4.07256 | 4.40E-58 |
| 22. | STM14_RS11405 | Propanediol dehydratase large subunit PduC | 4.06915 | 2.03E-23 |
| 23. | STM14_RS23400 | Ornithine carbamoyltransferase | 4.00766 | 6.76E-20 |
| 24. | STM14_RS10065 | Manganese efflux pump MntP | 3.83616 | 8.09E-21 |
| 25. | STM14_RS20040 | Acetolactate synthase 1 small subunit | 3.83222 | 4.58E-55 |
| 26. | STM14_RS11430 | Propanediol utilization microcompartment protein PduA | 3.76332 | 1.18E-05 |
| 27. | STM14_RS20130 | Cytochrome c-type biogenesis protein CcmH | 3.76324 | 1.53E-18 |
| 28. | STM14_RS10070 | 23S rRNA (guanine(745)-N(1))-methyltransferase | 3.72569 | 1.786E-46 |
| 29. | STM14_RS23405 | Carbamate kinase | 3.71075 | 3.365E-16 |
| 30. | STM14_RS12275 | Bifunctional murein DD-endopeptidase/murein LD-carboxypeptidase | 3.6602 | 2.284E-68 |

**Supplemental-table 4.** Differentially regulated biofilm genes upon treatment with 15μM clarithromycin.

| **S. No** | **Gene ID** | **Product** | **Log 2**<br>**(Treated**<br>**Vs**<br>**Control)** | **P- Value** |
| --- | --- | --- | --- | --- |
| **1** | STM14_RS06210 | Transcriptional regulator CsgD | -0.9831 | 3.25E-05 |
| **2** | STM14_RS06205 | Curli production assembly/transport protein CsgE | -1.1489 | 1.73E-09 |
| **3** | STM14_RS06215 | Curli minor subunit CsgB | -0.7701 | 0.0118 |
| **4** | STM14_RS06220 | Major curlin subunit CsgA | -0.6679 | 0.01857 |
| **5** | STM14_RS09410 | Cyclic di-GMP phosphodiesterase | -0.508 | 0.0243 |
| **6** | STM14_RS14685 | Biofilm-associated protein BapA | -2.0857 | 7.59E-11 |
| **7** | STM14_RS06195 | Curli production assembly/transport protein CsgG | -0.2800 | 0.1756 |
| **8** | STM14_RS06200 | Curli production assembly/transport protein CsgF | -0.9982 | 4.67E-07 |
| **9** | STM14_RS06225 | Curli assembly protein CsgC | 0.1310 | 0.7075 |
| **10** | STM14_RS19170 | Cellulosebiosynthesisprotein BcsF | 2.174 | 3.46E-09 |
| **11** | STM14_RS19155 | Cellulosebiosynthesisprotein BcsQ | 0.930 | 1.87E-07 |
| **12** | STM14_RS19140 | Cellulase | 0.1735 | 0.3603 |
| **13** | STM14_RS19160 | Hypotheticalprotein | -0.120 | 0.603 |
| **14** | STM14_RS19150 | UDP-formingcellulosesynthasecatalyticsubunit | -0.515 | 0.002 |
| **15** | STM14_RS19165 | Cellulosebiosynthesisprotein BcsE | -0.648 | 0.0002 |
| **16** | STM14_RS19175 | Cellulosebiosynthesisprotein BcsG | -0.735 | 0.00068 |
| **17** | STM14_RS19145 | Cellulosebiosynthesiscyclicdi-GMP-bindingregulatoryprotein BcsB | -1.053 | 7.86E-07 |
| **18** | STM14_RS19135 | Cellulosebiosynthesisprotein BcsC | -1.339 | 3.96E-11 |
| **19** | STM14_RS03355 | Fimbriae_biosynthesis_transcriptional_regulator_FimZ | 1.1946 | 0.0010 |
| **20** | STM14_RS03360 | Fimbriae_biosynthesis_regulator_FimY | 1.1304 | 3.34E-05 |
| **21** | STM14_RS02605 | Phosphate_response_regulator_transcription_factor_phob | 0.1914 | 0.2752 |
| **22** | STM14_RS02610 | Phosphate_regulon_sensor_histidine_kinase_phor | 0.0743 | 0.727 |
| **23** | STM14_RS15415 | Transcriptional_regulator_HilD | 0.6058 | 0.0435 |
| **24** | STM14_RS15420 | Transcriptional_regulator_HilA | -0.335 | 0.3021 |
| **25** | STM14_3659 | Fimbrial_protein | -2.214 | 6.94E-07 |
| **26** | STM14_5493 | Fimbrial_protein | 3.018 | 7.06E-07 |
| **27** | STM14_0234 | Fimbrial_major_subunit_StfA | 2.520 | 5.00E-06 |

**Supplemental-table 5.** Differentially regulated flagellar regulon genes upon treatment with 15μM clarithromycin.

| **S. No** | **Gene ID** | **Product** | **Log 2**<br>**(Treated**<br>**Vs**<br>**Control)** | **P- Value** |
| --- | --- | --- | --- | --- |
| 1 | STM14_RS06390 | flagellar_basal_body_rod_protein_FlgF | 2.998764 | 4.97E-14 |
| 2 | STM14_RS06375 | flagellar_basal_body_rod_protein_FlgC | 2.89797 | 6.48E-09 |
| 3 | STM14_RS06400 | flagellar_basal_body_L-ring_protein_FlgH | 2.52822 | 2.47E-10 |
| 4 | STM14_RS06370 | flagellar_basal_body_rod_protein_FlgB | 2.4967 | 1.29E-05 |
| 5 | STM14_RS10720 | flagella_biosynthesis_regulatory_protein_FliT | 2.3134 | 9.84E-08 |
| 6 | STM14_RS10715 | flagellar_export_chaperone_FliS | 2.2964 | 7.15E-08 |
| 7 | STM14_RS06420 | flagellar_hook-filament_junction_protein_FlgL | 2.1086 | 2.31E-09 |
| 8 | STM14_RS10525 | flagellar_motor_stator_protein_MotA | 2.0872 | 2.28E-15 |
| 9 | STM14_RS10530 | flagellar_transcriptional_regulator_FlhC | 2.0504 | 1.97E-21 |
| 10 | STM14_RS06385 | flagellar_hook_protein_FlgE | 1.8460 | 4.30E-07 |
| 11 | STM14_RS06395 | flagellar_basal_body_rod_protein_FlgG | 1.8250 | 3.46E-06 |
| 12 | STM14_RS06365 | flagellar_basal_body_P-ring_formation_protein_FlgA | 1.8200 | 1.47E-06 |
| 13 | STM14_RS10520 | motility_protein_MotB | 1.7858 | 6.81E-09 |
| 14 | STM14_RS10510 | chemotaxis_protein_CheW | 1.7074 | 8.89E-08 |
| 15 | STM14_RS06410 | flagellar_assembly_peptidoglycan_hydrolase_FlgJ | 1.6327 | 2.67E-05 |
| 16 | STM14_RS06355 | flagella_biosynthesis_chaperone_FlgN | 1.6005 | 7.78E-14 |
| 17 | STM14_RS10765 | flagellar_assembly_protein_FliH | 1.5823 | 0.00233 |
| 18 | STM14_RS10515 | chemotaxis_protein_CheA | 1.5598 | 1.71E-10 |
| 19 | STM14_RS06380 | flagellar_hook_assembly_protein_FlgD | 1.5537 | 9.17E-05 |
| 20 | STM14_RS10535 | flagellar_transcriptional_regulator_FlhD | 1.4339 | 1.90E-10 |
| 21 | STM14_RS06405 | flagellar_basal_body_P-ring_protein_FlgI | 1.3667 | 2.69E-05 |
| 22 | STM14_RS09880 | flagellar_brake_protein_YcgR | 1.3292 | 2.70E-05 |
| 23 | STM14_RS10710 | flagellar_filament_capping_protein_FliD | 1.293 | 4.61E-07 |
| 24 | STM14_RS00625 | molecular_chaperone_DnaK | 1.1719 | 1.38E-08 |
| 25 | STM14_RS06415 | flagellar_hook-associated_protein_FlgK | 1.1451 | 3.02E-06 |
| 26 | STM14_RS10730 | lipoprotein | 1.0210 | 1.88E-07 |
| 27 | STM14_RS10775 | flagella_biosynthesis_chaperone_FliJ | 1.0123 | 0.16042 |
| 28 | STM14_RS10785 | flagellar_basal_body-associated_protein_FliL | 0.9842 | 0.0227 |
| 29 | STM14_RS10750 | flagellar_hook-basal_body_complex_protein_FliE | 0.9788 | 0.16780 |
| 30 | STM14_RS10790 | flagellar_motor_switch_protein_FliM | 0.9611 | 0.00365 |
| 31 | STM14_RS10500 | protein-glutamate_O-methyltransferase_CheR | 0.7281 | 0.0737 |
| 32 | STM14_RS10485 | protein_phosphatase_CheZ | 0.6777 | 0.0249 |
| 33 | STM14_RS10755 | flagellar_basal_body_M-ring_protein_FliF | 0.6323 | 0.0266 |
| 34 | STM14_RS10760 | flagellar_motor_switch_protein_FliG | 0.61323 | 0.09108 |
| 35 | STM14_RS10480 | flagellar_type_III_secretion_system_protein_FlhB | 0.5606 | 0.34846 |
| 36 | STM14_RS10770 | flagellum-specific_ATP_synthase_FliI | 0.5440 | 0.348295 |
| 37 | STM14_RS10495 | chemotaxis_response_regulator_protein-glutamate_methylesterase | 0.5341 | 0.28957 |
| 38 | STM14_RS10780 | flagellar_hook_length_control_protein_FliK | 0.37276 | 0.384958 |
| 39 | STM14_RS10735 | selenium_metabolism_membrane_protein_YedE/FdhT | 0.3333 | 0.26309 |
| 40 | STM14_RS10800 | flagellar_type_III_secretion_system_protein_FliO | 0.2367 | 0.74662 |
| 41 | STM14_RS10490 | two-component_system_response_regulator | 0.1620 | 0.588553 |
| 42 | STM14_RS10545 | universal_stress_protein_UspC | 0.15765 | 0.33933 |
| 43 | STM14_RS10740 | putative_sulfurtransferase_YedF | 0.03712 | 0.9264 |
| 44 | STM14_RS10805 | flagellar_type_III_secretion_system_pore_protein_FliP | -0.024 | 0.95800 |
| 45 | STM14_RS10795 | flagellar_motor_switch_protein_FliN | -0.212 | 0.69754 |
| 46 | STM14_RS10505 | methyl-accepting_chemotaxis_protein_II | -0.423 | 0.209112 |
| 47 | STM14_RS06360 | anti-sigma-28_factor_FlgM | -0.4307 | 0.05950 |
| 48 | STM14_RS10815 | flagellar_type_III_secretion_system_protein_FliR | -0.509 | 0.17478 |
| 49 | STM14_RS10475 | flagellar_biosynthesis_protein_FlhA | -0.539 | 0.15473 |
| 50 | STM14_RS10810 | flagellar_biosynthesis_protein_FliQ | -0.559 | 0.34358 |
| **51** | STM14_RS07580 | anti-FlhDC_factor_YdiV | -1.223 | 1.13E-10 |
| **52** | STM14_RS10705 | flagellin_FliC | -1.2365 | 1.36E-08 |
| **53** | STM14_RS10820 | transcriptional_regulator_RcsA | -1.2452 | 0.00073 |
| **54** | STM14_RS10470 | flagellar_protein_FlhE | -1.7035 | 8.61E-06 |
| **55** | STM14_RS10725 | alpha-amylase | -2.2750 | 7.52E-31 |
| **56** | STM14_RS10690 | RNA_polymerase_sigma_factor_FliA | 2.2852 | 2.00E-16 |
| **57** | STM14_RS14875 | FliC/FljB_family_flagellin | -0.834 | 1.31E-06 |

**Supplemental table 6.**
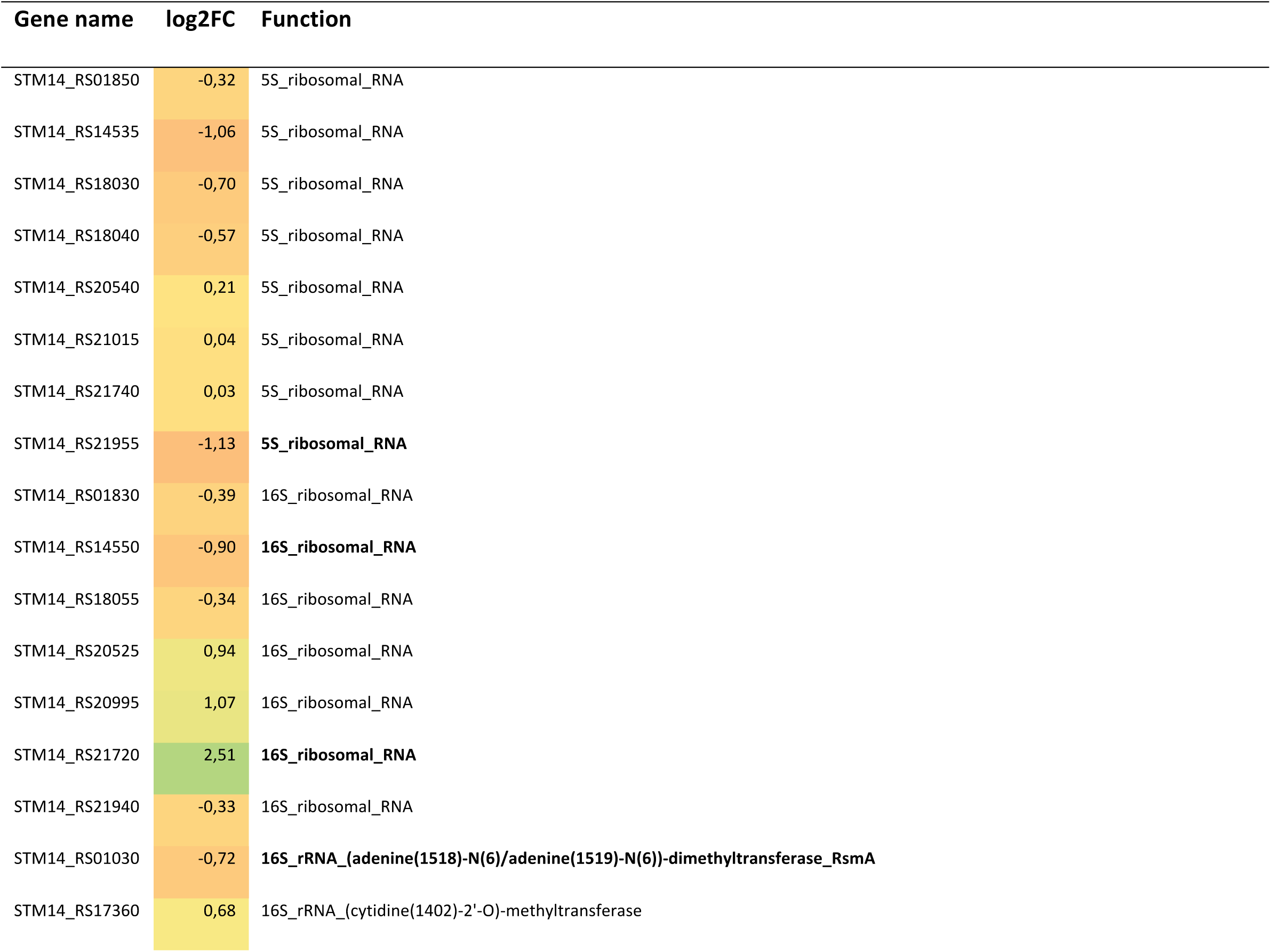
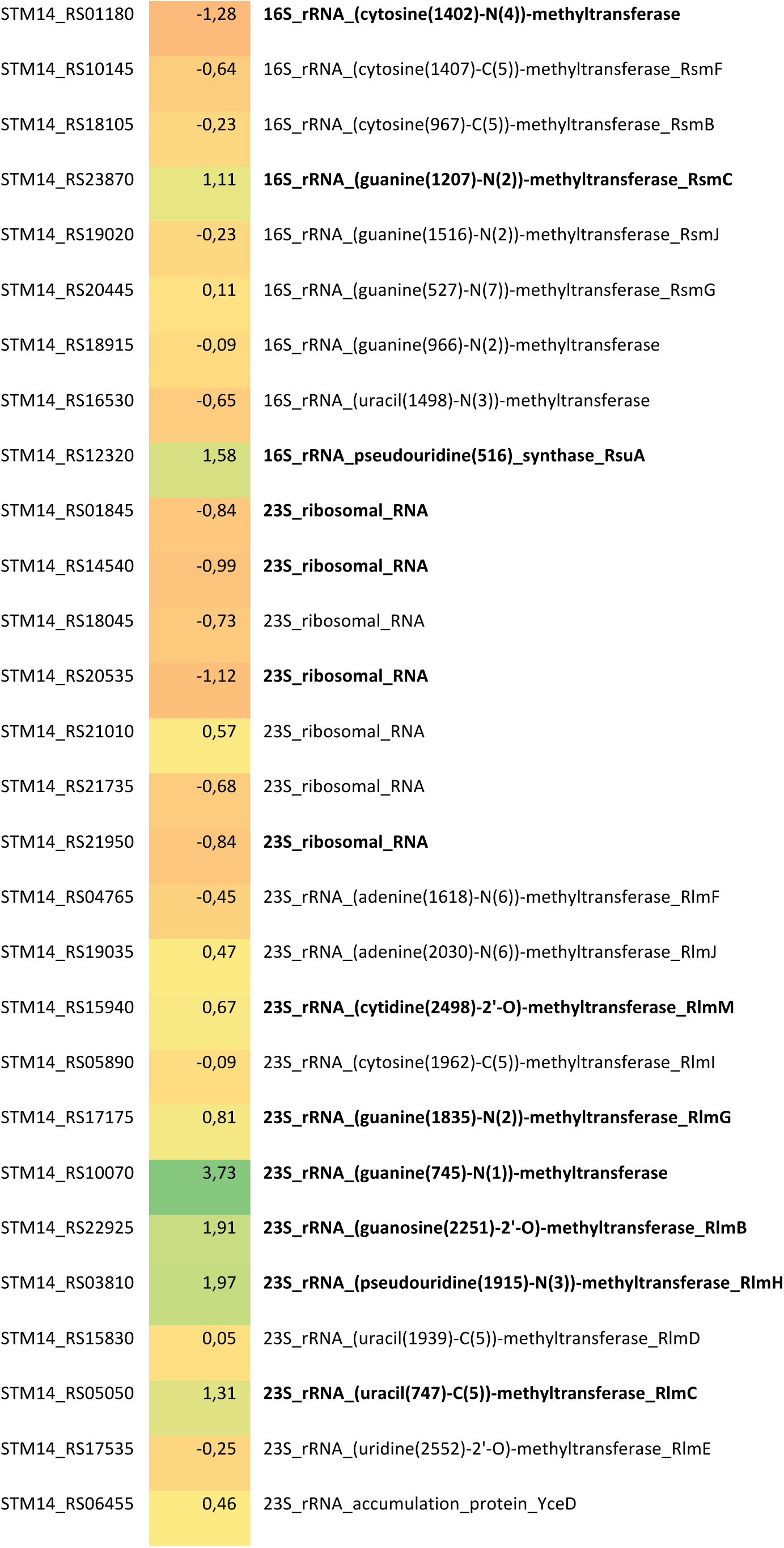
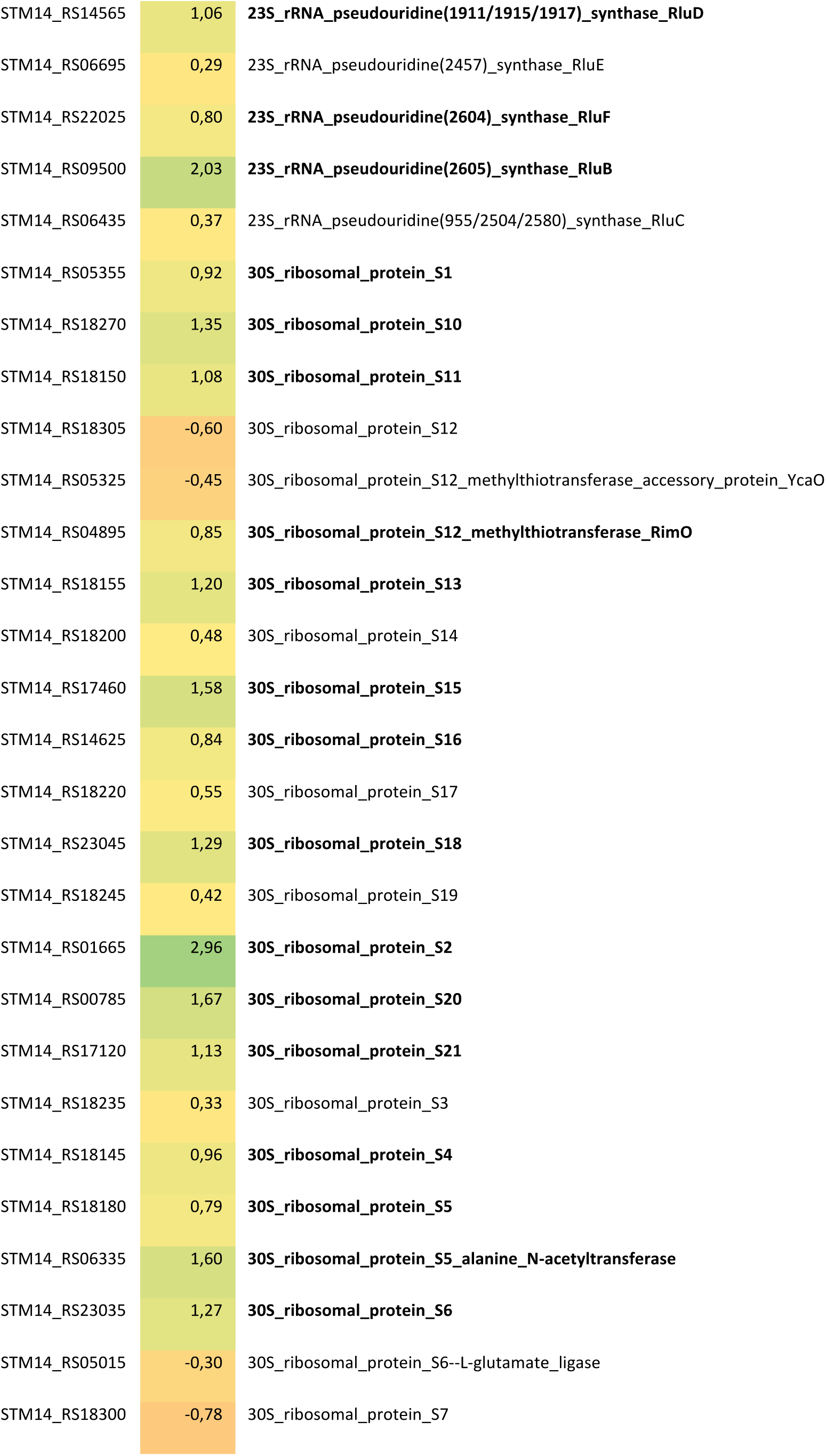
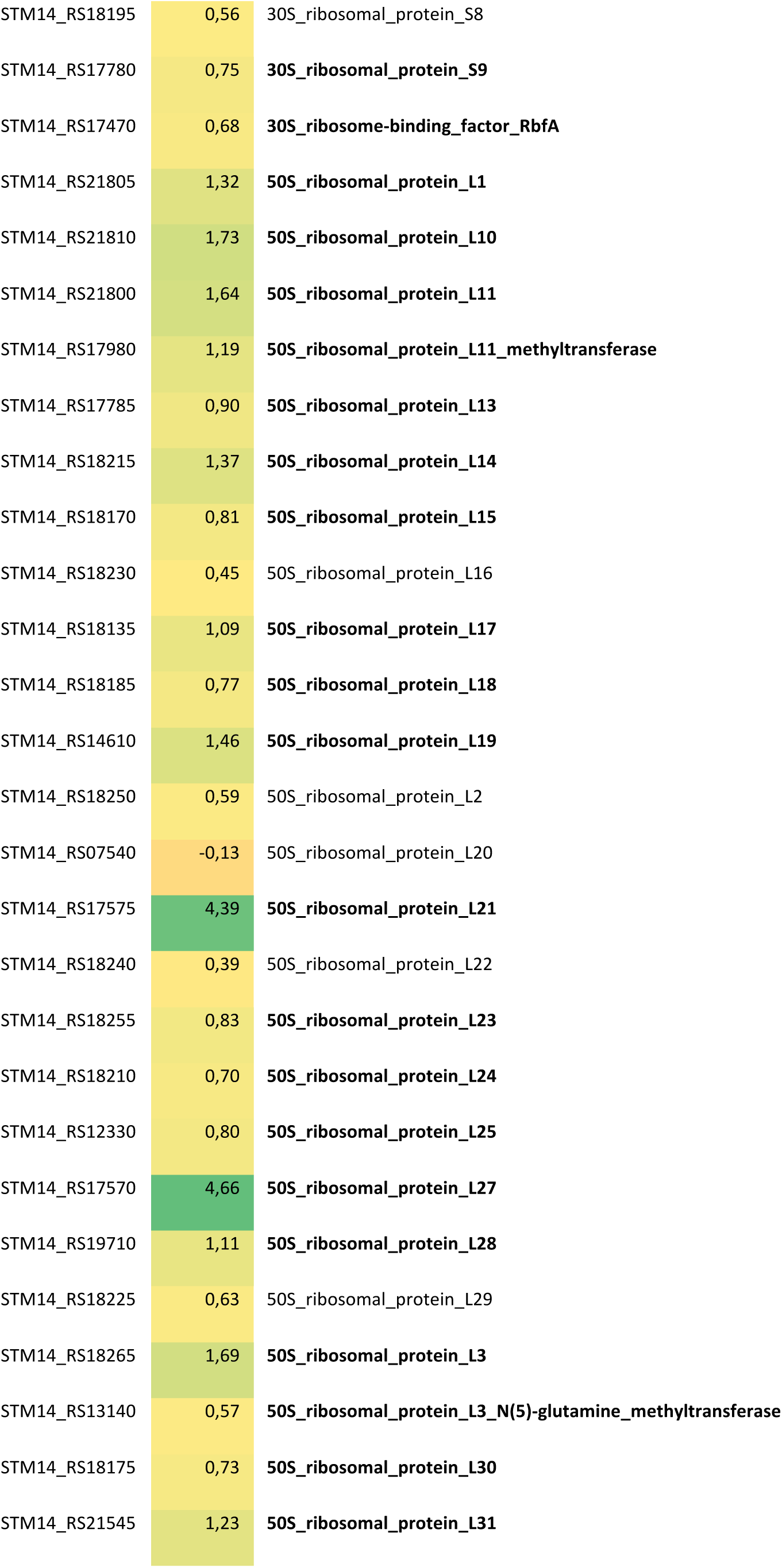
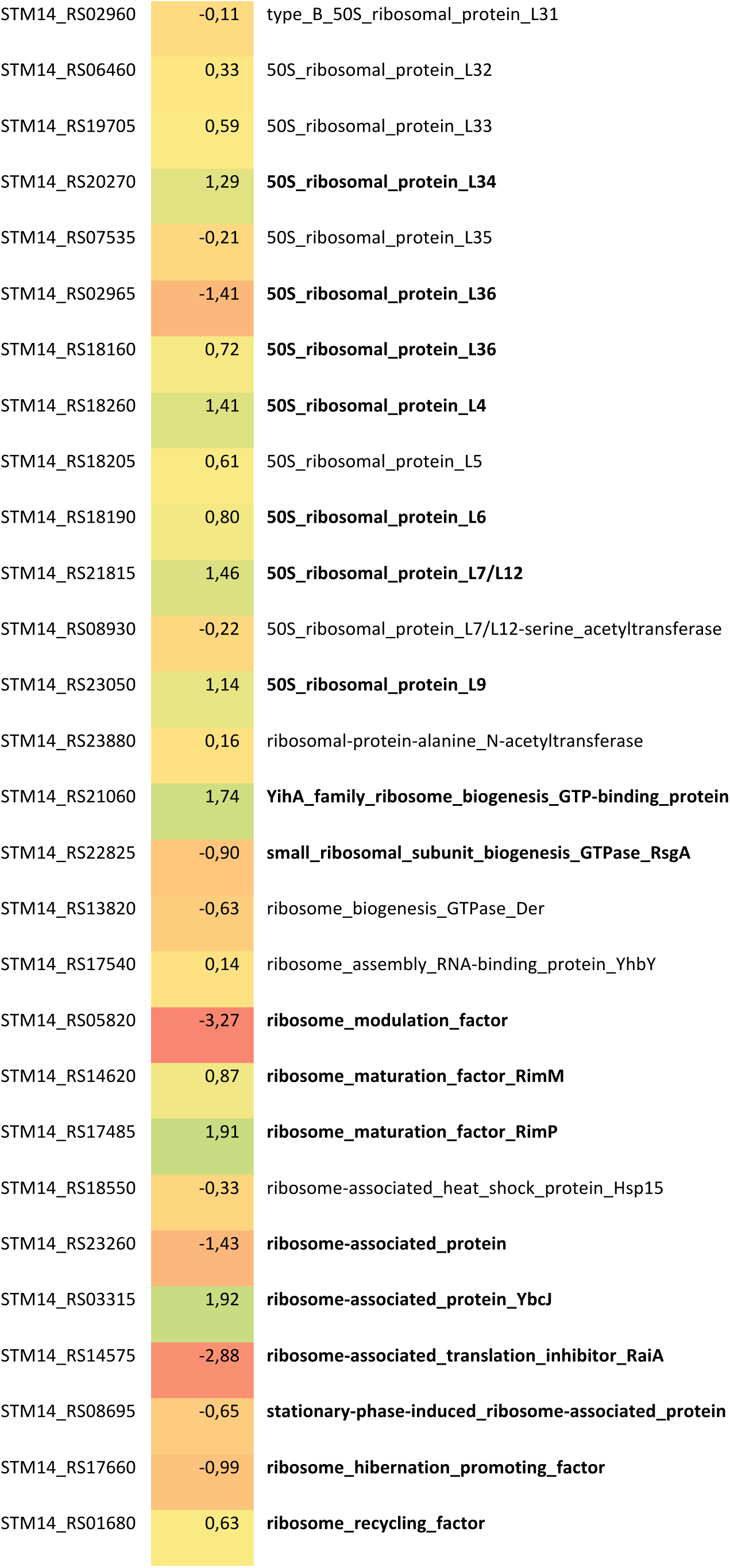
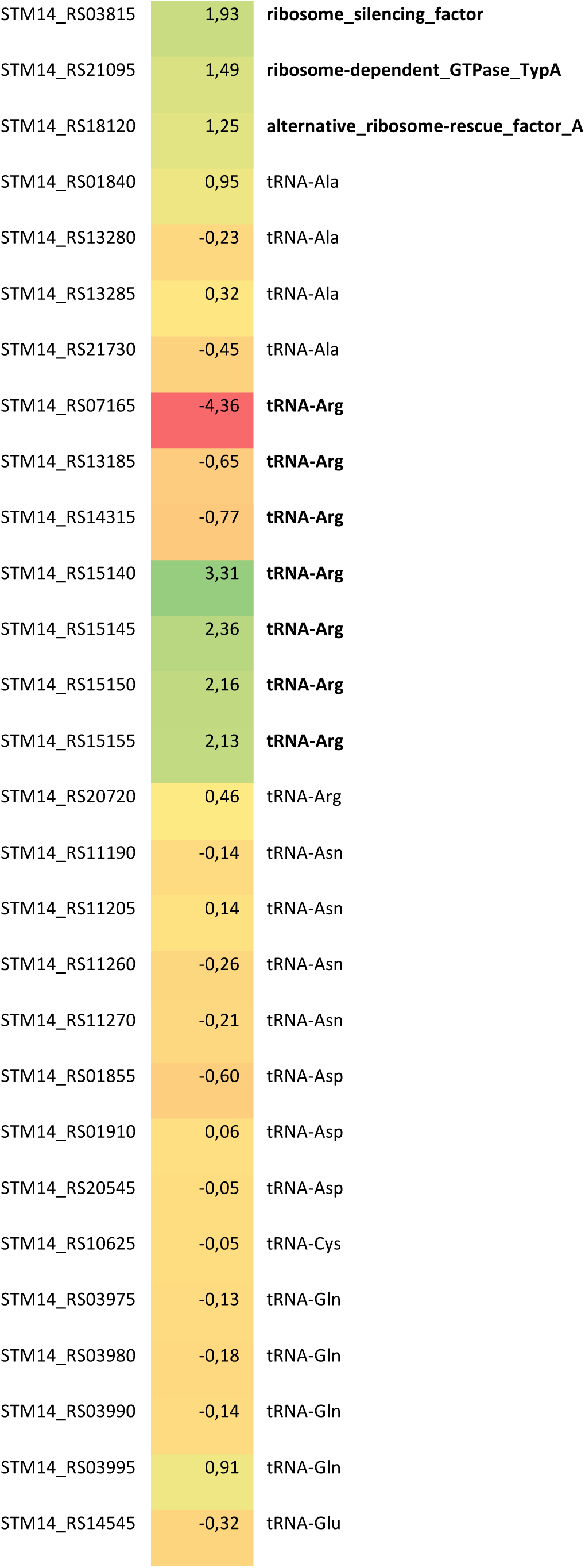
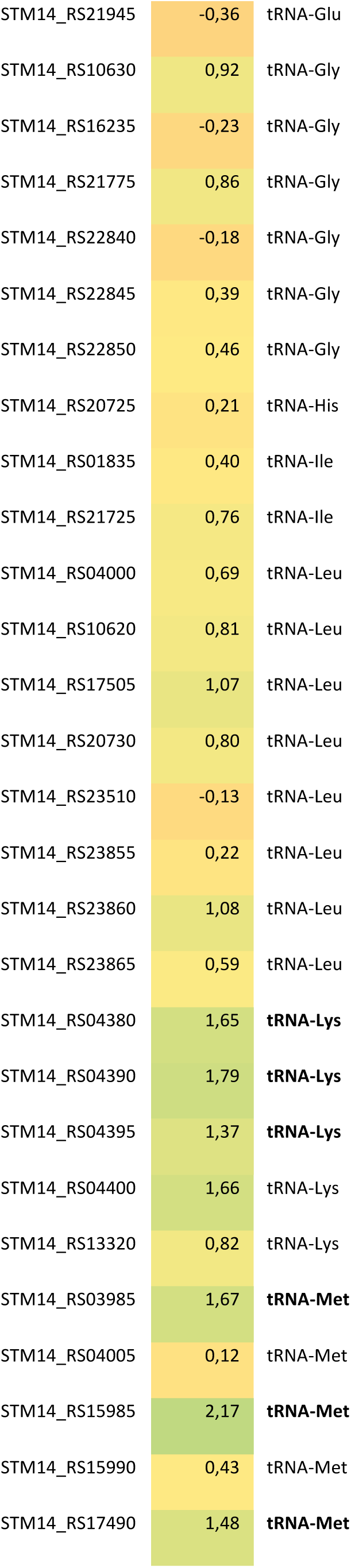
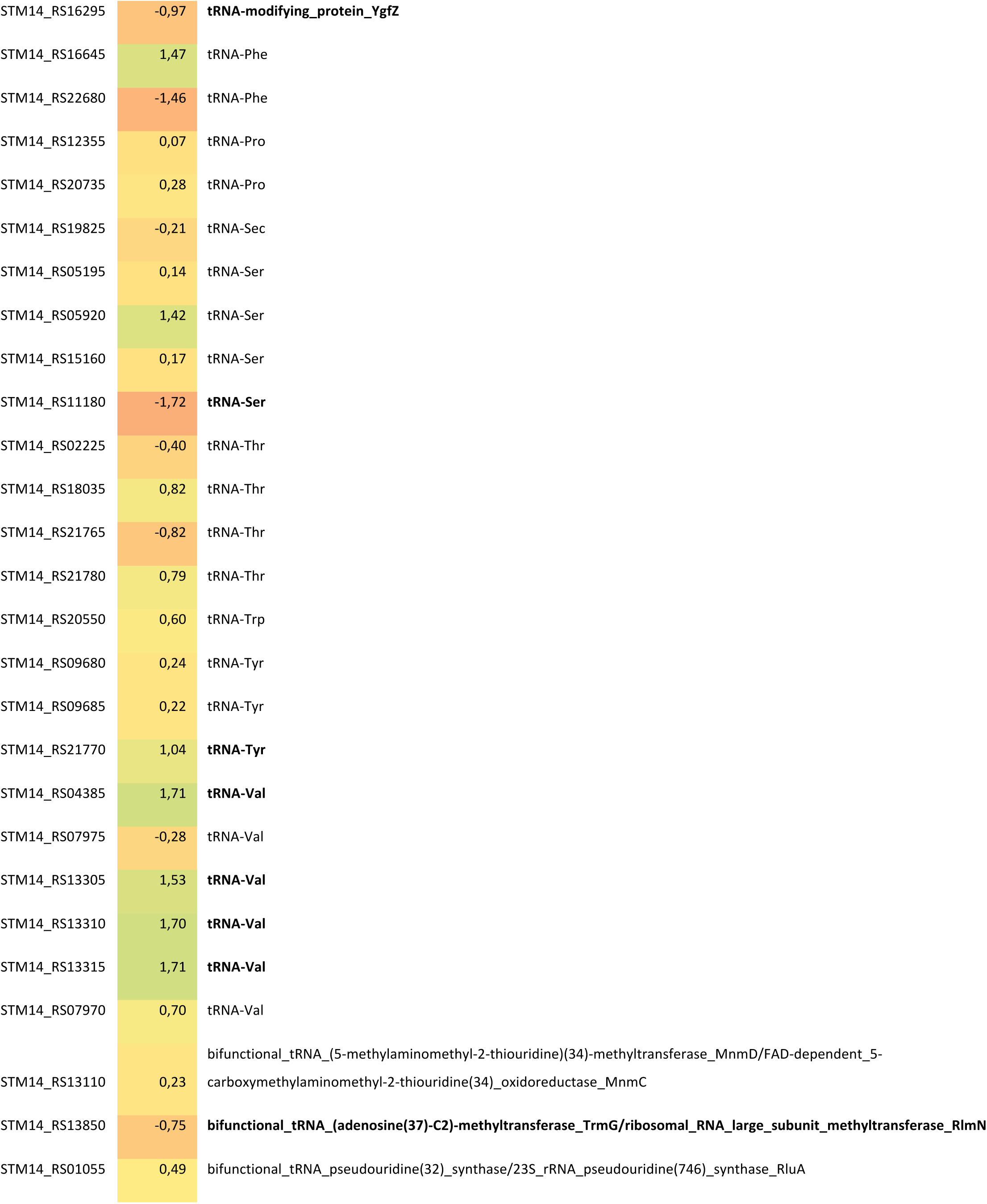
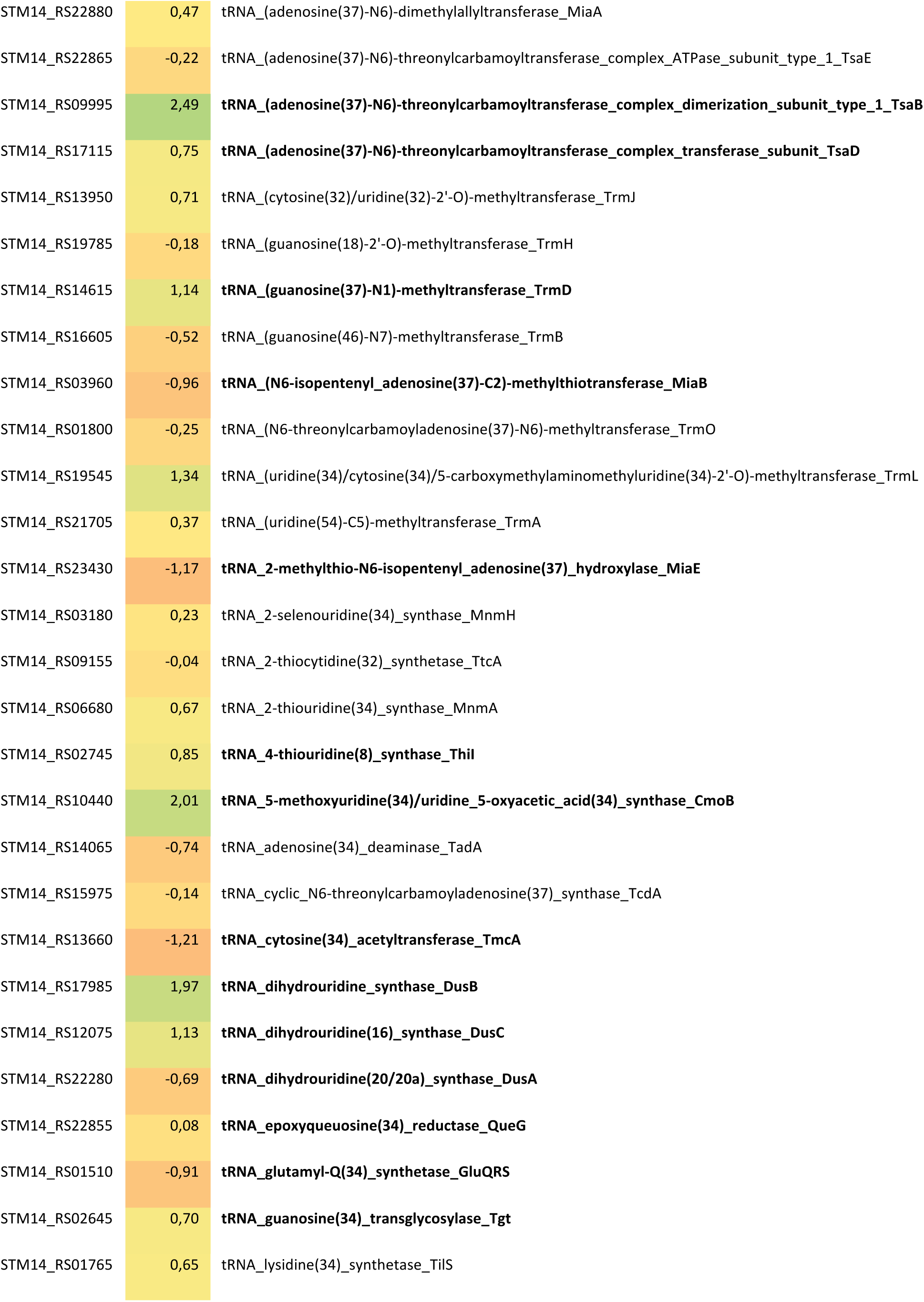
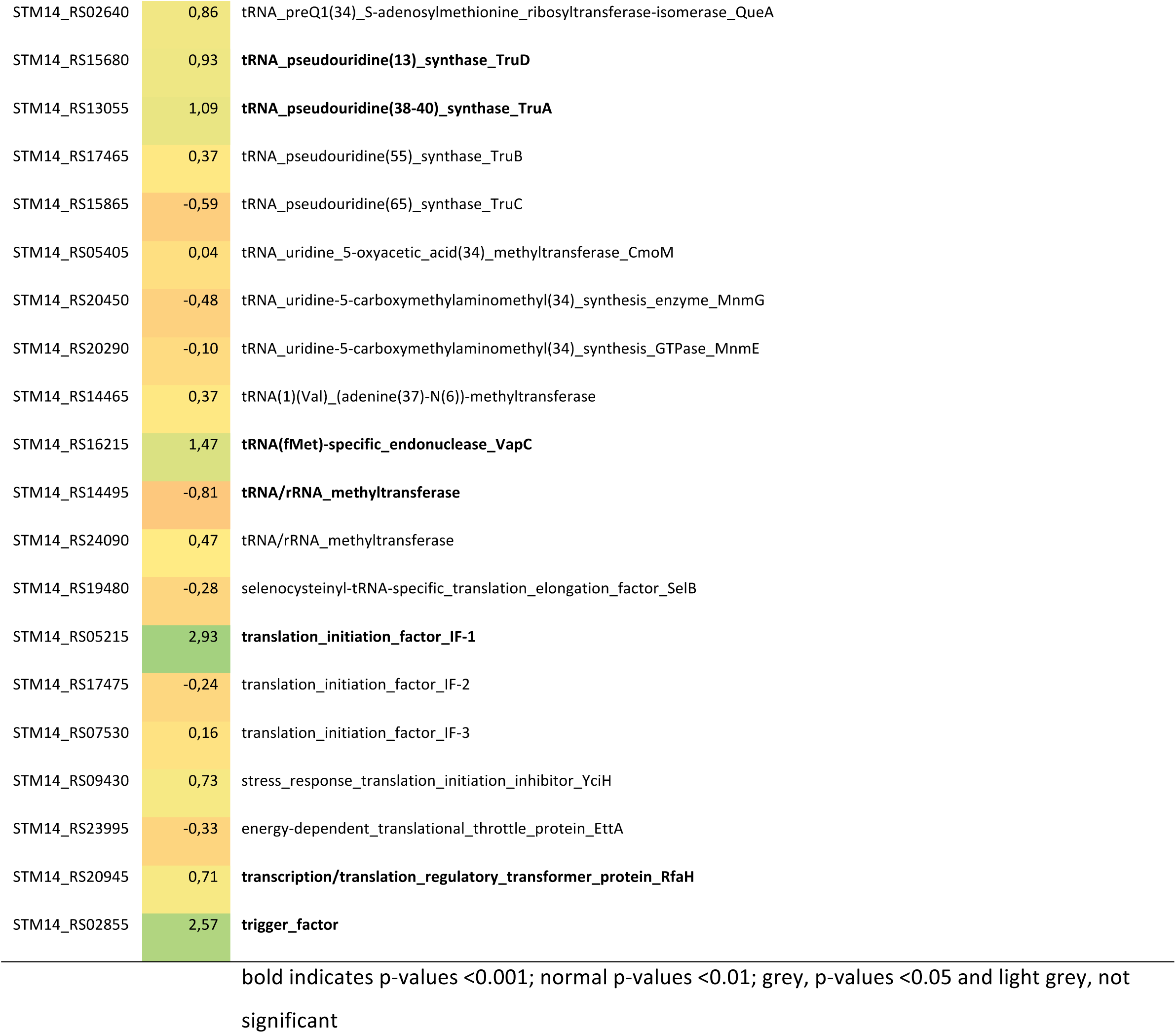
Heatmap of selection of differentially regulated ribosome and translation associated genes.

